# A novel sialic acid-binding adhesin present in multiple species contributes to the pathogenesis of Infective endocarditis

**DOI:** 10.1101/2020.07.17.206995

**Authors:** Meztlli O. Gaytán, Anirudh K. Singh, Shireen A. Woodiga, Surina A. Patel, Seon-Sook An, Arturo Vera-Ponce de León, Sean McGrath, Anthony R. Miller, Jocelyn M. Bush, Mark van der Linden, Vincent Magrini, Richard K. Wilson, Todd Kitten, Samantha J. King

## Abstract

Bacterial binding to platelets is a key step in the development of infective endocarditis (IE). Sialic acid, a common terminal carbohydrate on host glycans, is the major receptor for streptococci on platelets. So far, all defined interactions between streptococci and sialic acid on platelets are mediated by serine rich repeat proteins (SRRPs). However, we identified *Streptococcus oralis* subsp. *oralis* IE-isolates that bind sialic acid but lack SRRPs. In addition to binding sialic acid, some SRRP-negative isolates also bind the cryptic receptor β-1,4-linked galactose through a yet unknown mechanism. Using comparative genomics, we identified a novel sialic acid-binding adhesin, here named AsaA (<u>a</u>ssociated with <u>s</u>ialic acid <u>a</u>dhesion A), present in IE-isolates lacking SRRPs. We demonstrated that *S. oralis* subsp. *oralis* AsaA is required for binding to platelets in a sialic acid-dependent manner. AsaA comprises a non-repeat region (NRR), consisting of a FIVAR/CBM and two Siglec-like and Unique domains, followed by 31 DUF1542 domains. When recombinantly expressed, Siglec-like and Unique domains competitively inhibited binding of *S. oralis* subsp. *oralis* and directly interacted with sialic acid on platelets. We further demonstrated that AsaA impacts the pathogenesis of *S. oralis* subsp. *oralis* in a rabbit model of IE. Additionally, we found AsaA orthologues in other IE-causing species and demonstrated that the NRR of AsaA from *Gemella haemolysans* blocked binding of *S. oralis* subsp. *oralis*, suggesting that AsaA contributes to the pathogenesis of multiple IE-causing species. Finally, our findings provide evidence that sialic acid is a key factor for bacterial-platelets interactions in a broader range of species than previously appreciated, highlighting its potential as a therapeutic target.

**Authors summary:** Infective endocarditis (IE) is typically a bacterial infection of the heart valves that causes high mortality. Infective endocarditis can affect people with preexisting lesions on their heart valves (Subacute-IE). These lesions contain platelets and other host factors to which bacteria can bind. Growth of bacteria and accumulation of host factors results in heart failure. Therefore, the ability of bacteria to bind platelets is key to the development of IE. Here, we identified a novel bacterial protein, AsaA, which helps bacteria bind to platelets and contributes to the development of disease. Although this virulence factor was characterized in *Streptococcus oralis*, a leading cause of IE, we demonstrated that AsaA is also present in several other IE-causing bacterial species and is likely relevant to their ability to cause disease. We showed that AsaA binds to sialic acid, a terminal sugar present on platelets, thereby demonstrating that sialic acid serves as a receptor for a wider range of IE-causing bacteria than previously appreciated, highlighting its potential as a therapeutic target.

## Introduction

The human oral cavity is inhabited by more than 700 bacterial species. *Streptococcus oralis, Streptococcus mitis* and *Gemella haemolysans* are among the early colonizers of the oral cavity [1, 2]. Although often associated with oral health, oral commensals can also gain access to the bloodstream where some can cause diseases including infective endocarditis (IE) [3-5].

IE is typically a bacterial infection of the heart valve endothelium. One of the hallmarks of IE is the formation of vegetations, produced by the accumulation of host factors and bacterial proliferation. These vegetations can affect heart valve function and lead to congestive heart failure [6]. IE can have two presentations. Acute IE is sudden and severe; it typically affects previously normal heart valves and is commonly caused by staphylococci. Subacute IE develops gradually and has a more subtle onset, it affects previously damaged heart valves and is commonly caused by oral streptococci, including *S. oralis* [7].

Although the exact mechanisms that lead to IE development are unknown, it is proposed that bacterial binding to platelets plays a key role in the pathogenesis of this disease [8]. For many streptococcal species, binding to platelets involves the direct interaction of serine rich repeat proteins (SRRPs) with sialic acid, a common terminal carbohydrate [9-15]. SRRPs are large, surface-associated glycoproteins that form fibrils. Members of this adhesin family possess a modular organization, typically including an N-terminal secretion signal that mediates export through an accessory Sec system, and a C-terminal cell-wall anchoring domain. SRRPs also have a non-repeat region (NRR) that mediates adhesion and one or two regions consisting of serine-rich repeats (SRR),which are highly glycosylated and proposed to serve as stalks that extend the NRR from the cell surface [16-18].

The NRR of many sialic acid-binding SRRPs consists of a sialic acid binding immunoglobulin-like lectin (Siglec)-like and a Unique domain [9-11, 15, 19, 20]. The Unique domain is proposed to influence the conformation of the Siglec-like domain [21], which directly interacts with sialic acid. The Siglec-like domains of the SRRPs Fap1 (*S. oralis*), GspB (*Streptococcus gordonii*), Hsa (*S. gordonii*) and SrpA (*Streptococcus sanguinis*) contain an arginine residue essential for sialic acid binding within the semi-conserved YTRY motif [9-11, 15, 21, 22].

Studies have demonstrated that the expression of some SRRPs containing Siglec-like and Unique domains is important in IE [15, 23-25]. Since the arginine residue, essential for binding of the SRRP GspB to sialic acid on platelets, contributes to the pathogenesis of *S. gordonii* in a rat model of IE, virulence is attributed to the specific interaction of SRRPs with sialic acid [15]. Hence, the current paradigm establishes that binding to sialic acid on platelets via Siglec-like and Unique domain-containing SRRPs contributes to the pathogenesis of IE.

The SRRP-sialic acid binding mechanism has been described in the IE-causing bacteria *S. gordonii, S. sanguinis* and *S. oralis* [9-11, 15, 19, 20]. However, many other bacterial species cause IE through yet undefined mechanisms. Recent work demonstrated that, in addition to binding sialic acid on platelets, *S. oralis* and *S. gordonii* also bind to β-1,4-linked galactose, exposed upon sialic acid removal [9, 26]. In *S. oralis*, adhesion to both sialic acid and β-1,4-linked galactose requires the SRRP Fap1 [9]. This novel strategy of binding multiple receptors reveals that streptococcal-platelet interactions are more complex than previously considered.

In this study, we identified *S. oralis* subsp. *oralis* IE-isolates that bind sialic acid and β-1,4-linked galactose but lack SRRPs. We demonstrated that these strains have a novel sialic acid-binding adhesin, here named AsaA (<u>a</u>ssociated with <u>s</u>ialic acid <u>a</u>dhesion A), that contributes to the colonization of vegetations in an animal model of IE. AsaA orthologues were also found in several other IE-causing bacterial species suggesting this mechanism of adhesion contributes to the pathogenesis of multiple species.

## Results

### Serine-rich repeat proteins are not essential for adhesion of *S. oralis* subsp. *oralis* to sialic acid

Bacterial binding to platelets is a key step in the development of IE [8]. Many streptococcal species, including *S. oralis* subsp. *oralis*, bind to sialic acid on platelets via an SRRP [9-11, 15, 19, 20]. Here we demonstrated that only two of five *S. oralis* subsp. *oralis* IE-isolates screened encode the sialic acid-binding SRRP Fap1 (Fig 1A). Since previous reports established that Fap1 binds sialic acid, we tested adhesion of the IE-isolates to this carbohydrate by competing binding with a carbohydrate-binding module (CBM) specific for sialic acid (CBM40, [27]). As shown in Fig 1B, adhesion of the five IE-isolates tested was significantly reduced by CBM40. This indicates that while sialic acid is a conserved receptor for all strains tested, Fap1 is not essential for binding this carbohydrate.

**Fig 1.**
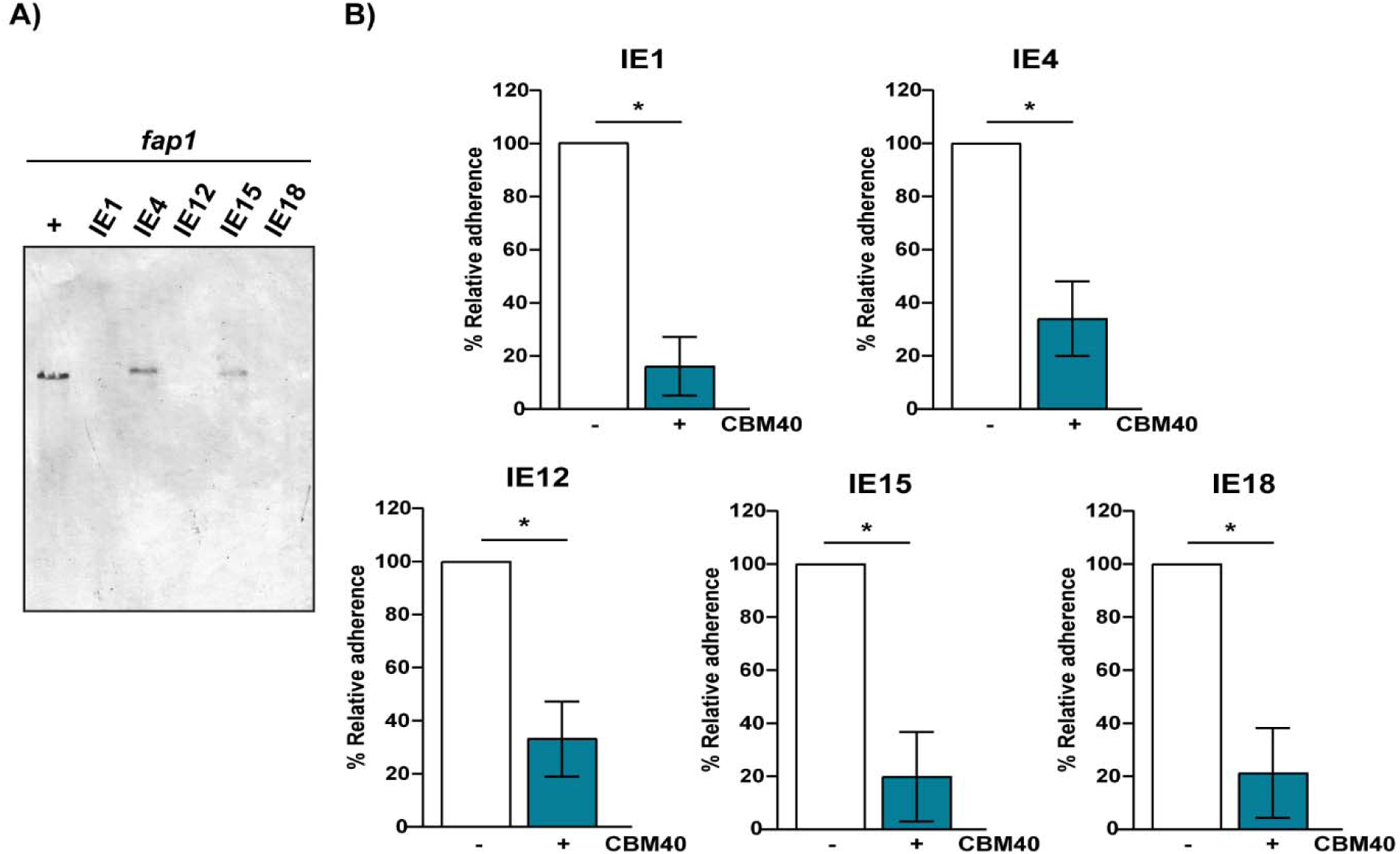
Fap1 is not essential for adhesion to sialic acid. (A) Southern blot of EcoRV-digested gDNA from five different *S. oralis* subsp. *oralis* IE-isolates using a DIG-labeled *fap1* probe. gDNA from *S. oralis* subsp. *oralis* strain ATCC10557 was used as a positive control (+). *fap1* was detected in only two IE-isolates, (B) Adherence of five *S. oralis* subsp. *oralis* IE-isolates to platelets in the presence of 30 µM CBM40 (+). Adherence was expressed as a percentage relative to the same strain in the absence of CBM40 (-). Values are the means for at least three independent experiments, each performed in triplicate, ± SD. *Statistical significance was tested by two-tailed Student’s t-test. *, P ≤ 0.0015.

Isolates that bind sialic acid but lack *fap1* must either encode a distinct SRRP or employ an alternative adhesion mechanism. To address the first possibility, we screened for the presence of the three different *secA2* variants identified in *S. oralis* (Uo5, F0392 and ATCC6249). SecA2 is the ATPase that energizes export of SRRPs and is more highly conserved than members of the SRRP family [28]. The gene encoding SecA2 was only detected in the *fap1*^+^ isolates, suggesting that the *fap1*^*-*^ isolates lack other SRRPs (S1 Fig). The absence of genes encoding an SRRP and the machinery required for its export were confirmed by genomic sequencing of two *fap1*^-^ isolates. Overall, these data indicate that sialic acid is a conserved receptor for *S. oralis* subsp. *oralis*, but that SRRPs are not essential for binding this carbohydrate. Instead, SRRP^-^ isolates must bind sialic acid through a novel mechanism.

### β-1,4-linked galactose can serve as a receptor for some SRRP^-^ *S. oralis* subsp. *oralis* isolates

We previously demonstrated that β-1,4-linked galactose, exposed upon removal of sialic acid by bacterial neuraminidase, serves as an additional receptor for *S. oralis* subsp. *oralis* binding to platelets [9]. Binding to β-1,4-linked galactose was reported to be Fap1 dependent, hence, it was unclear whether the *fap1*^-^ isolates can bind this carbohydrate. In order to address this question, carbohydrates underlying sialic acid were exposed by neuraminidase pretreatment and binding to β-1,4-linked galactose tested by determining whether CBMs previously shown to specifically bind this carbohydrate (CBM71-1.2) can competitively inhibit adhesion [29]. As previously reported for other Fap1^+^ isolates, the two Fap1^+^ strains included in this study were significantly reduced in adhesion upon neuraminidase treatment and further reduced upon addition of CBM71-1.2 (Fig 2B). This result supports previous data demonstrating that Fap1^+^ isolates bind both sialic acid and β-1,4-linked galactose [9]. Binding of the *fap1*^*-*^ isolates IE1 and IE18 was not reduced by removal of sialic acid, indicating that binding to underlying carbohydrates is as efficient as binding to sialic acid (Fig. 2A). The fact that addition of CBM71-1.2 significantly reduced adhesion of these strains to neuraminidase treated platelets demonstrates that these SRRP^-^ isolates can bind β-1,4-linked galactose. Unlike IE1 and IE18, binding of the SRRP^-^ isolate IE12 to neuraminidase treated platelets was not reduced by CBM71-1.2, indicating that this strain binds sialic acid but not β-1,4-linked galactose (Fig 2A). Together, our data demonstrate that SRRPs are not essential for binding to sialic acid and β-1,4-linked galactose. Therefore, SRRP^*-*^ strains must use an alternative mechanism to bind these carbohydrates. Furthermore, the differences in carbohydrate binding capabilities displayed by the SRRP^-^ isolates suggest that distinct adhesins mediate binding to sialic acid and β-1,4-linked galactose.

**Fig 2.**
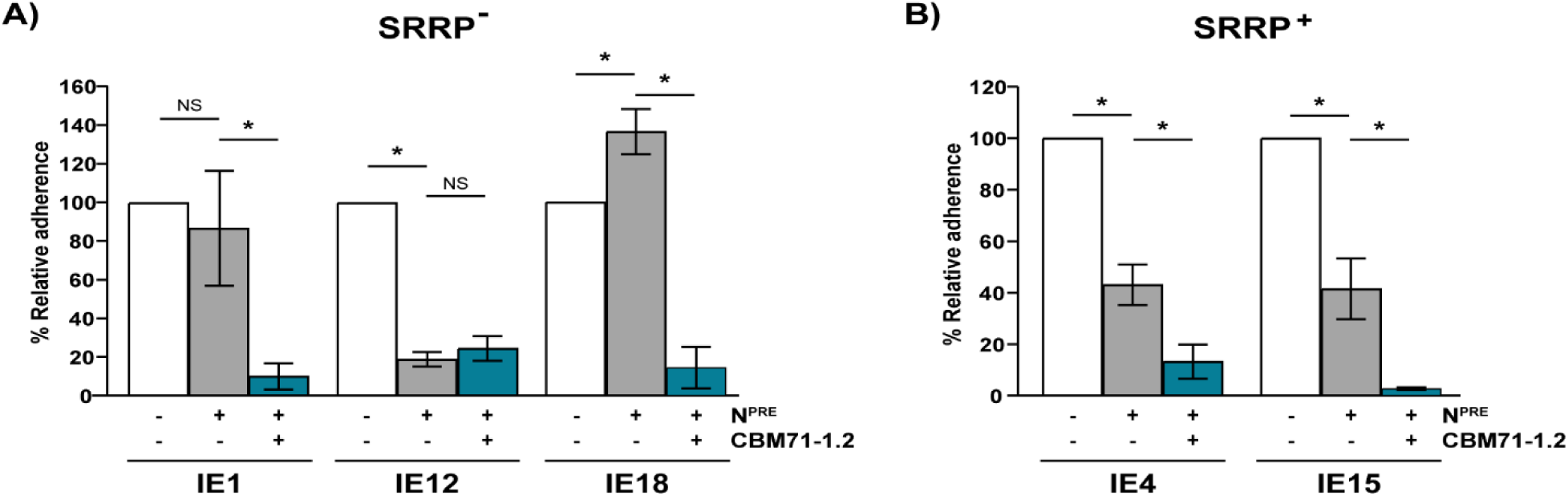
β-1,4-linked galactose serves as a receptor for some *S. oralis* subsp. *Oralis* isolates. Binding to β-1,4-linked galactose was tested by the addition of 50 µM CBM71-1.2 following pretreatment with neuraminidase (N^PRE^) or PBS (-). The graphs show adherence of the SRRP^-^ (A) and SRRP^+^ (B) IE-isolates to untreated or neuraminidase-pretreated platelets in presence or absence of CBM71-1.2. Adherence is expressed as a percentage relative to binding of the strains to untreated platelets in the absence of CBM71-1.2. Values are the means for at least three independent experiments, each performed in triplicate, ± SD. Statistical significance was tested by two-tailed Student’s t-test. *, P ≤ 0.015; NS, not significant.

### SRRP^-^ isolates employ a novel mechanism of adherence that requires a Sortase A-dependent surface protein(s)

To identify the mechanism(s) by which *S. oralis* subsp. *oralis* strains lacking SRRPs bind to carbohydrates on platelets, we first tested the effect of a Sortase A (SrtA) mutant on bacterial binding. SrtA mediates covalent attachment of many adhesins to the cell wall of Gram-positive bacteria [30, 31]. Deletion of *srtA* from SRRP^-^ isolates, IE12 and IE18, significantly reduced bacterial binding to platelets (Fig 3A). Complementation of the *srtA* mutants significantly increased adhesion, confirming that the phenotype observed was due to the mutation introduced. These results indicate that a surface protein(s), attached to the cell wall by SrtA, is required for adhesion. Sialic acid serves as the main receptor for IE12 and IE18 on platelets (Fig 1B). However, IE18 also binds β-1,4-linked galactose exposed upon sialic acid removal (Fig 2A). Adhesion of an IE18 *srtA* mutant to neuraminidase-pretreated platelets was also reduced (Fig 3A). The adhesion phenotype was recovered upon complementation. These results indicate that a SrtA-dependent surface protein(s) is also required for adhesion to β-1,4-linked galactose.

**Fig 3.**
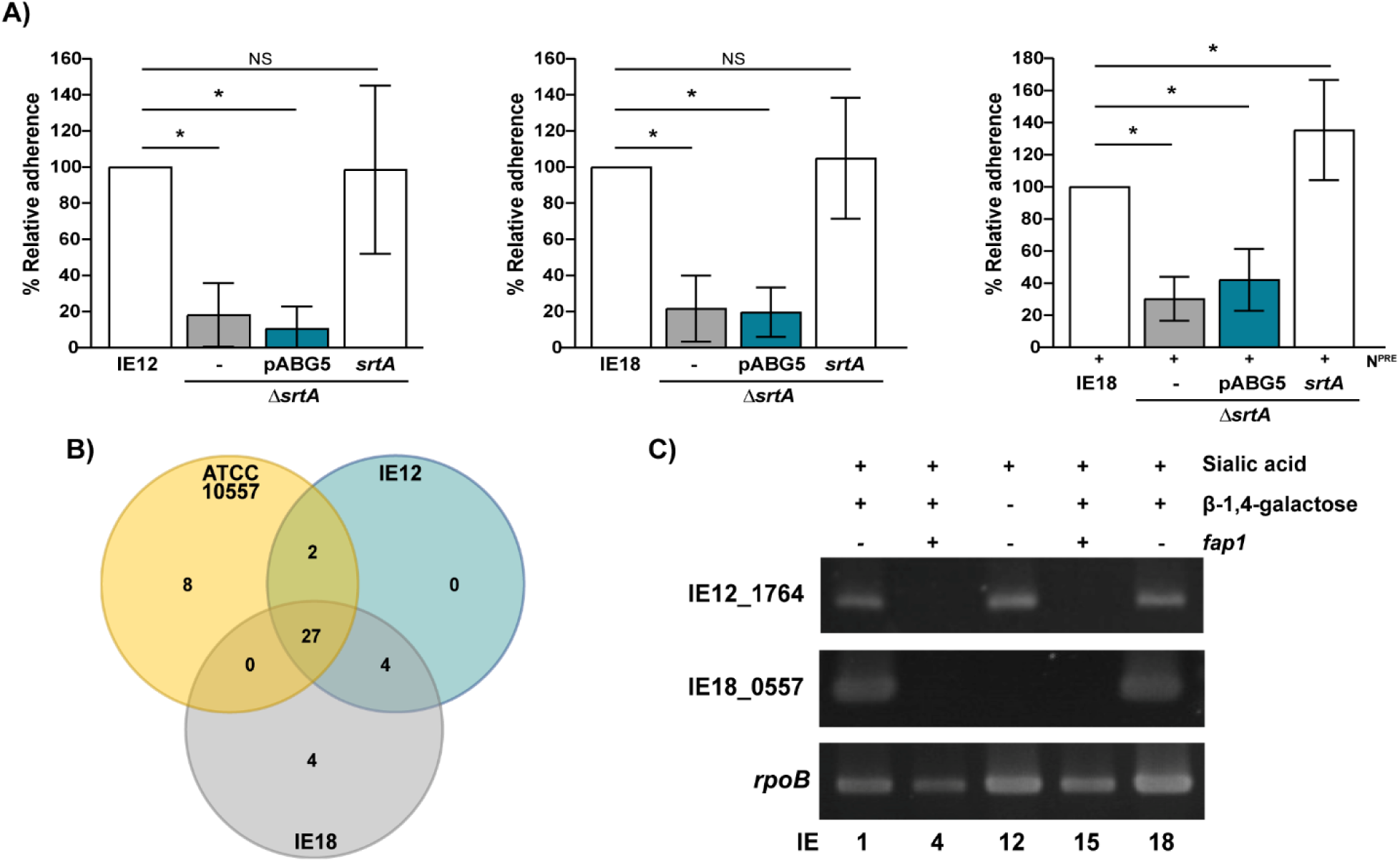
A Sortase A-dependent surface protein(s) is required for binding of *S. oralis* IE-isolates lacking SRRPs. (A) a *srtA* mutant is reduced in binding to platelets. Graphs indicate adherence of IE12 and IE18 *srtA* mutants and complements, with *srtA* or the empty vector (pABG5), relative to parental strains. The contribution of SrtA-dependent surface proteins to β-1,4-linked galactose binding was tested by comparing adhesion to neuraminidase-pretreated platelets (N^PRE^). (B) Venn diagram showing shared and unique LPxTG-containing proteins in three different *S. oralis* subsp. *oralis* IE-isolates. Comparisons were made by BLASTP, proteins with >80% amino acid identity were considered as shared between strains. (C) As demonstrated by PCR, the presence of IE12_1764 and IE18_0557 correlates with the ability of *fap1*^-^ isolates to bind sialic acid or β-1,4-linked galactose, respectively. *rpoB* was used as a positive control. Adherence data are the means for at least three independent experiments, each performed in triplicate, ± SD. Statistical significance was tested by two-tailed Student’s t-test. *, P ≤ 0.006.

The above experiments established that SRRP^-^ isolates bind to platelets through a novel mechanism that requires a surface protein(s), anchored by SrtA. To identify putative sialic acid and β-1,4-linked galactose binding adhesins we performed comparative genomic analysis. Since SrtA recognizes surface proteins that possess an LPxTG amino acid motif [32], the comparative analysis was restricted to genes encoding proteins with this motif. Potential sialic acid-binding adhesins were identified as LPxTG-containing proteins encoded by IE12 and IE18, two SRRP^-^ isolates that bind this carbohydrate, but absent in the SRRP^+^ strain ATCC10557. Proteins encoded by IE18, but absent in IE12 and ATCC10557, were considered as potential β-1,4-linked galactose binding adhesins. As seen in Fig 3B, four potential adhesins for each carbohydrate were identified (S1 Table). However, the distribution of only one potential adhesin for each carbohydrate correlated with the binding phenotypes of the isolates (Fig 3C and S2 Fig). These results support the hypothesis that SRRP^-^ isolates bind sialic acid and β-1,4-linked galactose through novel mechanisms involving the SrtA-dependent proteins encoded by IE12_1764 and IE18_0557, respectively.

### IE18_0557 encodes a Csh-like protein

IE18_0557 was the only gene encoding a putative β-1,4-linked galactose binding adhesin present in all SRRP^-^ strains that bind this carbohydrate (Fig 3C and S2 Fig). IE18_0557, which is expressed in IE18 (S3 Fig), is predicted to encode a 3097 amino acid protein that shares 76% amino acid identity with *S. gordonii* CshA (Sg_CshA). Sg_CshA is a fibril-forming adhesin involved in autoaggregation and adhesion to extracellular matrix proteins and other bacterial species [33-35]. The NRR of *S. gordonii* CshA consists of three distinct non-repetitive domains: NR1, NR2 and NR3. The NR2 adopts a lectin-like fold, which is characteristic of carbohydrate binding proteins [36]. In addition, the expression of CshA is associated with lactose-sensitive interactions between *S. gordonii* and *Actinomyces naeslundii* [33]. Together, these data suggest that the protein encoded by IE18_0557, which we have named “Csh-like,” is involved in binding to β-1,4-linked galactose.

The Csh-like protein encoded by IE18 contains an YSIRK type signal peptide, an NRR and 23 repeats of the CshA-type fibril repeat (Fig 4A). Homology detection and structural prediction using HHpred [37], revealed that the NRR of this protein shares structural similarity with the NR2 of *S. gordonii* CshA (PDB: 5L2D_B, [36]), supporting the hypothesis that Csh-like is required for binding to β-1,4-linked galactose. To validate this hypothesis, we generated a *csh-like* mutant. However, when performing adherence assays with the IE18 *csh-like* mutant bacterial numbers added to the assay were inconsistent; after examination with Gram-staining we observed that the mutant was forming aggregates (S4 Fig) that were not disrupted by vortexing and hence interfered with the platelet-binding assays. As an alternative approach, we examined the effect of adding recombinant Csh-like NRR (Csh_NRR) to IE18 adhesion assays. As expected, binding of IE18 to neuraminidase-pretreated platelets was reduced by CBM71-1.2; however, the addition of Csh_NRR did not significantly reduce binding (Fig 4B). Proper folding of the Csh_NRR was confirmed by circular dichroism (CD) spectroscopy (S5 Fig). These results do not support the hypothesis that Csh-like mediates binding of SRRP^-^ strains to β-1,4-linked galactose.

**Fig 4.**
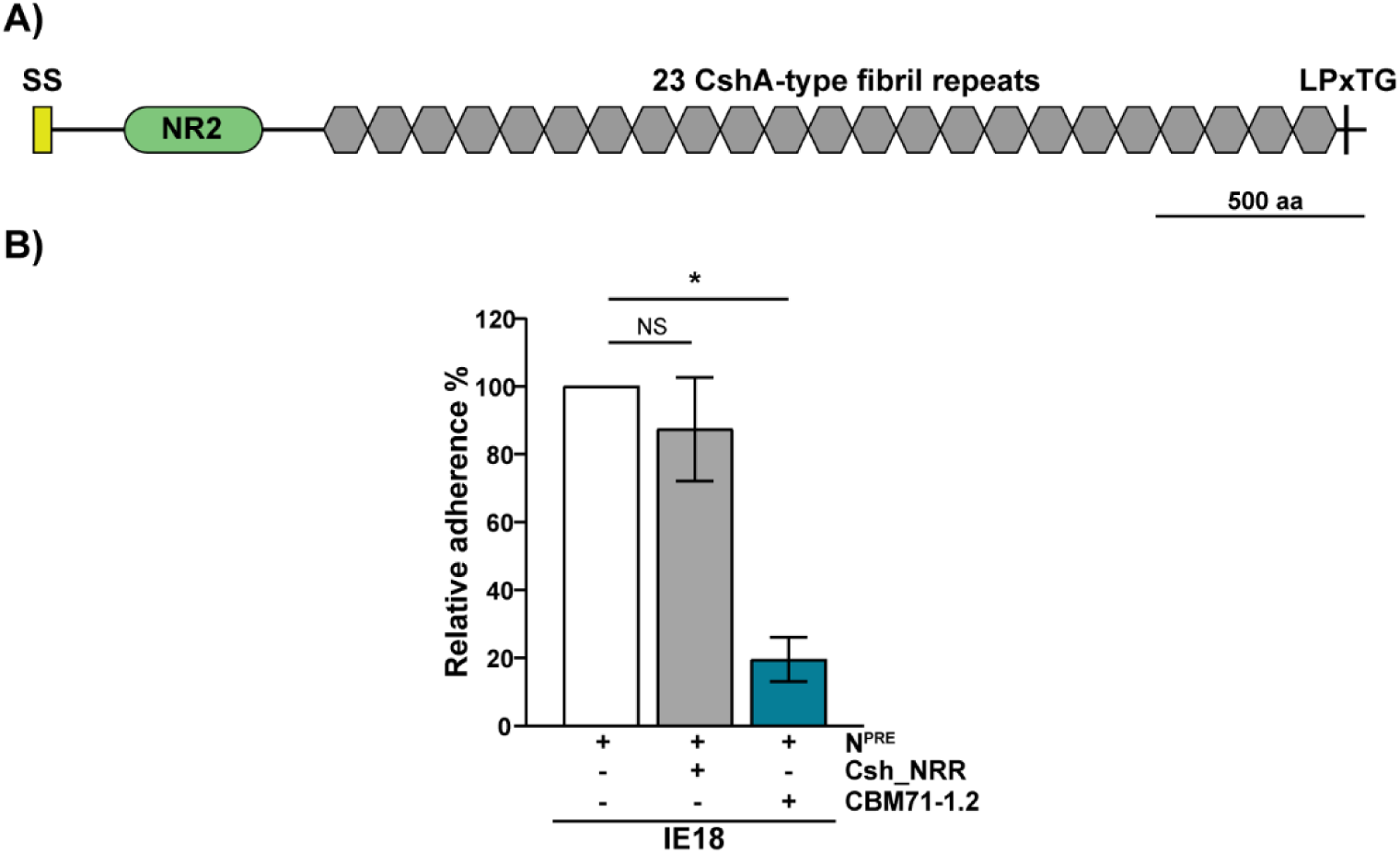
Csh-like protein is not involved in IE18 binding to β-1,4-linked galactose. (A) Schematic illustrating the predicted protein architecture of Csh-like from *S. oralis* subsp. *oralis* IE18 (IE18_0557). Csh-like consists of an N-terminal secretion signal (SS), followed by the NR2 domain (PDB: 5L2D), 23 CshA-type fibrils repeats and a cell wall anchoring motif (LPxTG). (B) The non-repeat region of Csh-like (Csh_NRR) is not involved in binding to β-1,4-linked galactose. Unlike CBM71-1.2, the addition of 30 µM of the recombinantly expressed Csh_NRR did not significantly reduce binding of IE18 to neuraminidase-pretreated (N^PRE^) platelets. Adherence is expressed as a percentage relative to binding of IE18 to neuraminidase-pretreated (N^PRE^) platelets in the absence of CBM71-1.2 or Csh_NRR. Values are the means for at least three independent experiments, each performed in triplicate, ± SD. Statistical significance was tested by two-tailed Student’s t-test. *, P ≤ 0.001; NS, not significant.

### IE12_1764 encodes a putative sialic acid-binding protein

Through comparative genomics we identified four potential sialic acid-binding adhesins present in IE12 and IE18 (Fig 3B). However, only IE12_1764 (IE18_0535) was absent in Fap1^+^ strains but present in all SRRP^-^ strains that bind sialic acid (Fig 3C and S2 Fig). This putative sialic binding adhesin encoded by IE12 and IE18 consists of an N-terminal YSIRK-type secretion signal, followed by an NRR and 31 or 28 DUF1542 domains, respectively. The NRRs of these proteins share 98.72% amino acid identity, suggesting that they both perform the same function. To facilitate data interpretation, further studies were performed only in IE12, which unlike IE18, binds only to sialic acid.

IE12_1764, which we confirmed is expressed in IE12 (S3 Fig), encodes a 3146 amino acid protein. Homology detection and structural prediction using HHpred [37] revealed that the N-terminal region of the NRR shares structural homology with both a putative CBM (77.74% probability, PDB: 4NUZ) and a FIVAR (found in various architectures) domain (60.11% probability, PDB: 6GV8). FIVAR domains are also found in some characterized adhesins like Ebh and Embp from *Staphylococcus aureus* and *Staphylococcus epidermidis*, respectively, and are involved in binding to fibronectin [38]. The FIVAR/CBM region is followed by two predicted Siglec-like and Unique domains that share structural similarity with similar domains found in the sialic acid-binding SRRPs Hsa (99.38 and 99.26% probability, PDB: 6EFC), SrpA (99.21 and 99.15% probability, PDB: 5KIQ) and GspB (99.06 and 98.97% probability, PDB: 3QC5) (Fig 5A and B); however, most characterized SRRPs have only one Siglec-like and Unique domain [15, 21, 39]. The NRR is followed by 31 DUF1542 domains, which were previously shown to form fibrils when overexpressed [40]. Overall, these data lead to the hypothesis that IE12_1764 encodes a novel sialic acid-binding adhesin, here named AsaA (<u>A</u>ssociated with <u>s</u>ialic acid <u>a</u>dhesion).

**Fig 5.**
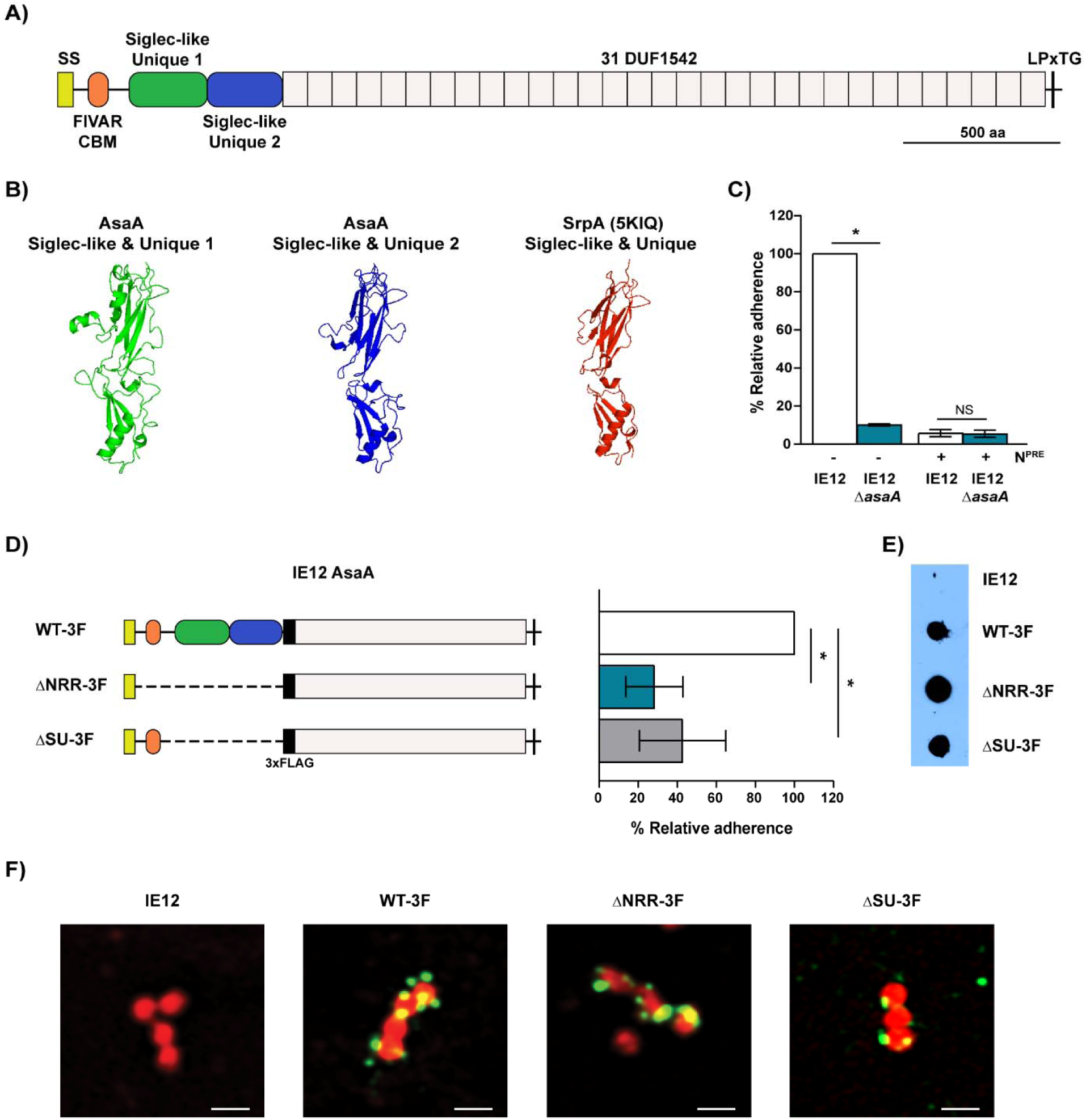
AsaA is a novel sialic acid-binding adhesin. (A) Schematic illustrating the predicted protein architecture of AsaA (IE12_1764) from *S. oralis* subsp. *oralis* IE12. AsaA consists of an N-terminal secretion signal (SS), followed by the non-repeat region predicted to contain a FIVAR/CBM domain and two Siglec-like and Unique domains, followed by 31 DUF1542 domains and a C-terminal LPxTG motif. (B) Comparison of the AsaA Siglec-like and Unique domains tertiary structure, predicted by SWISS-MODEL, to the solved structure of SrpA Siglec-like and Unique domain (5KIQ). (C) An IE12 *asaA* mutant was reduced in binding to platelets. Neuraminidase pretreatment (N^PRE^) did not further reduced binding of the mutant. (D) Left: schematic illustrating the domains removed in the isogenic IE12 variants. To further evaluate protein production and localization, a 3xFLAG tag (black box) was added to all constructs, including the parental strain (WT-3F). Right: Adhesion of WT-3F and the isogenic AsaA-deletion mutants to platelets. (E) Production of the AsaA deletion variants evaluated by the detection of the 3xFLAG tag in whole-cell lysates by immunodot-blot. (F) Detection of the AsaA-deletion variants on the cell surface by immunofluorescent microscopy. Representative images of Nile red-stained bacteria (red) incubated with anti-FLAG and Alexa Fluor 488-conjugated secondary antibody (green). Bars 1 µm. Adherence is expressed as a percentage relative to binding of the parental strain to untreated platelets. Values are the means for at least three independent experiments, each performed in triplicate, ± SD. Statistical significance was tested by two-tailed Student’s t-test. *, P ≤ 0.0004. NS, not significant.

### AsaA is a novel sialic acid-binding adhesin

A non-polar mutation in *asaA* (IE12 Δ*asaA*) significantly reduced binding of IE12 to platelets. Moreover, sialic acid removal by neuraminidase pretreatment did not further reduce adhesion of IE12 Δ*asaA* (Fig 5C), demonstrating that AsaA is required for bacterial binding to platelets in a sialic acid-dependent manner. Our attempts to complement the *asaA* mutant were unsuccessful, likely due to the high number of repeats; however, a second independent *asaA* mutant was similarly reduced in adhesion (S6 Fig).

AsaA consists of an NRR followed by 31 DUF1542 domains. In SRRPs, the NRR, specifically the Siglec-like and Unique domain within it, mediates adhesion to sialic acid. To dissect the role of the different AsaA domains in adhesion we constructed isogenic mutants lacking the entire NRR or the Siglec-like and Unique domains. To monitor protein stability and localization a 3xFLAG tag was added to the parental strain (IE12 *asaA* WT-3F) used to generate the mutants (IE12 *asaA* ΔNRR-3F and ΔSU-3F). Binding of the WT-3F strain to platelets was reduced as compared to the untagged IE12 (S7 Fig). However, the isogenic mutants lacking either the entire NRR or the Siglec-like and Unique domains were significantly reduced in adhesion to platelets as compared to the WT-3F parental strain (Fig 5D). Importantly, we demonstrated that the isogenic mutants are stable and localized on the cell surface (Fig 5E-F). These results indicate that the NRR of AsaA, specifically the Siglec-like and Unique domains, are required for bacterial binding to platelets.

Next, we demonstrated that the recombinantly expressed NRR (AsaA_NRR) competitively inhibits binding of IE12 to platelets in a dose-dependent manner (Fig 6A). IE12 adhesion was not reduced by glutathione *S*-transferase (GST) alone, indicating that the competitive inhibition of IE12 binding by AsaA_NRR is specific (S8 Fig). We confirmed that this reduction in adhesion is mediated by the predicted Siglec-like and Unique domains, since addition of a recombinant protein containing only these domains (AsaA_SU_1-2) significantly reduced binding. This reduction in binding is AsaA and sialic acid-dependent (Fig 6B).

**Fig 6.**
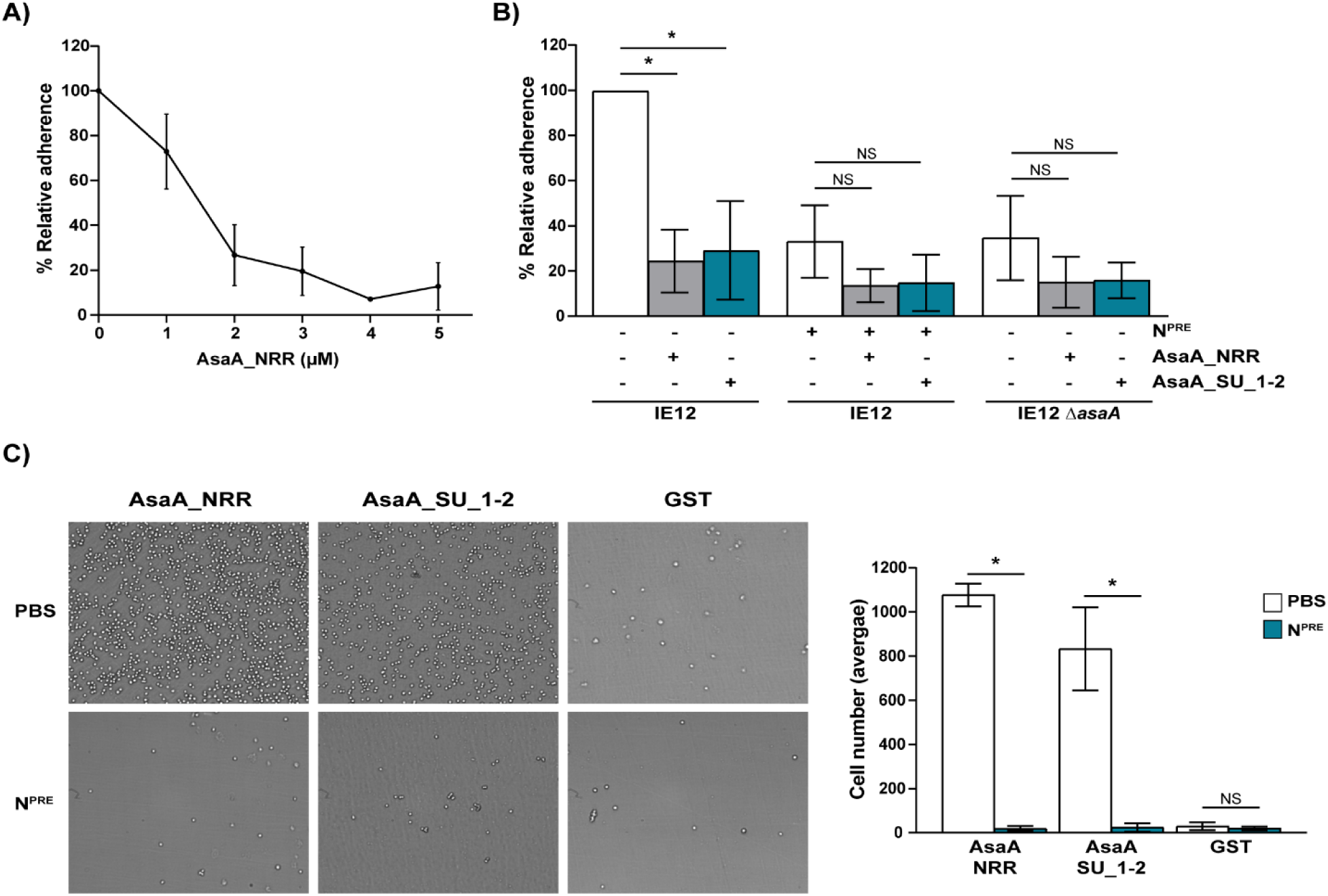
The Siglec-like and Unique domains of AsaA mediate direct binding to sialic acid on platelets. (A) Adhesion of IE12 was reduced in a dose-dependent manner by increasing concentrations of AsaA_NRR. (B) Adhesion of IE12 and the *asaA* mutant to untreated and neuraminidase pretreated (N^PRE^) platelets in the presence of 5 µM AsaA_NRR or AsaA_SU_1-2. Adherence is expressed as a percentage relative to binding of IE12 to untreated platelets in the absence of recombinant proteins. (C) Direct binding of platelets pretreated with PBS or neuraminidase (N^PRE^) to immobilized AsaA_NRR, AsaA_SU_1-2 or GST. A representative image of platelets binding to immobilized protein is shown. The graph shows the average number of platelets recovered in three independent experiments performed in triplicate. Statistical significance was tested by two-tailed Student’s t-test. *, P ≤ 0.002. NS, not significant.

Finally, we addressed whether the reduction in IE12 binding caused by the addition of AsaA_NRR and AsaA_SU_1-2 was due to the direct interaction of these recombinant proteins with sialic acid on platelets. To this end, we examined the ability of platelets pretreated with PBS or neuraminidase to bind to immobilized AsaA_NRR, AsaA_SU_1-2 or GST. While platelets pretreated with PBS bound efficiently to both recombinant proteins, essentially no binding of neuraminidase treated platelets was observed (Fig. 6C). Likewise, no binding of platelets to GST was observed. These results establish that AsaA directly binds sialic acid on platelets through the Siglec-like and Unique domains. In summary, our results demonstrate that *S. oralis* subsp. *oralis* AsaA is a novel adhesin that mediates direct binding to sialic acid on platelets.

### AsaA contributes to the colonization of vegetations by *S. oralis* IE12 in a rabbit model of IE

Since binding to sialic acid on platelets is a key step in the development of IE, the impact of AsaA on the virulence of IE12 was evaluated *in vivo*, using a rabbit model of IE [41]. To that end, the IE12 *asaA* mutant and a chloramphenicol-resistant derivative of IE12 (IE12 Cm^r^) were co-inoculated into rabbits in which aortic valve damage was induced by catheterization. The following day, rabbits were euthanized and vegetations in or near the aortic valves were removed. The mean colony-forming units (CFUs) recovered for the *asaA* mutant were significantly lower than those recovered for IE12 Cm^r^ (1.06×10^8^ vs 3.36×10^8^) (Fig 7). These studies demonstrate that *S. oralis* subsp. *oralis* AsaA contributes to the colonization of vegetations and is a potential virulence factor for IE.

**Fig 7.**
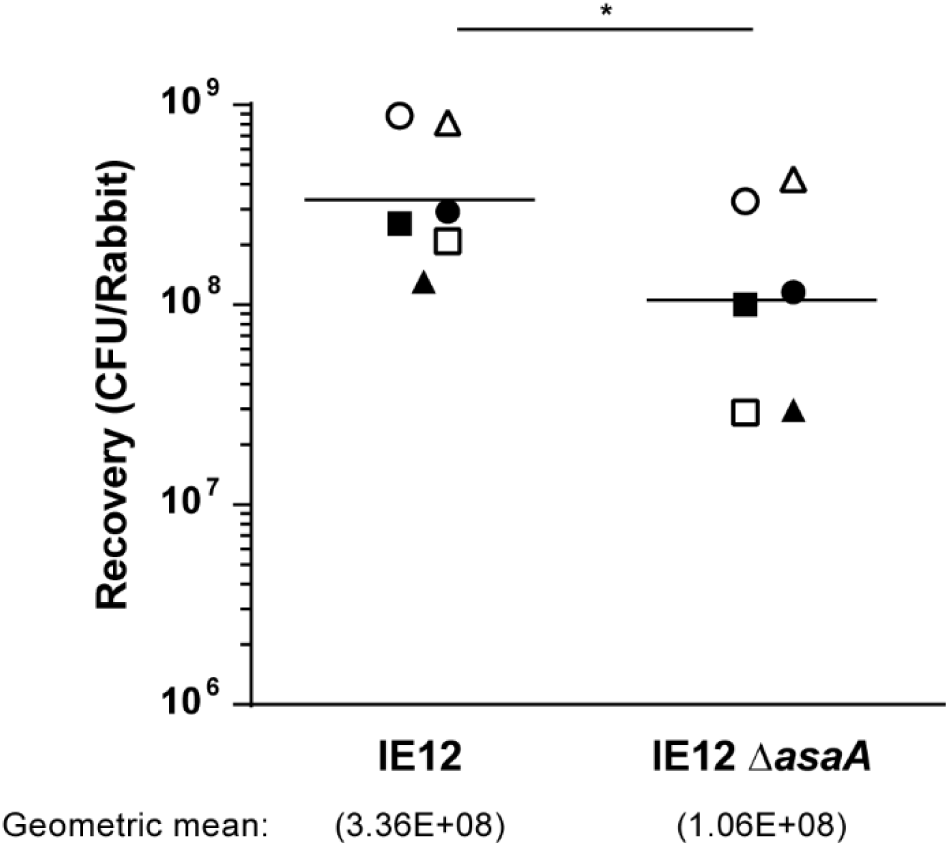
AsaA contributes to *S. oralis* IE12 virulence in a rabbit model of IE. The two strains indicated were co-inoculated into six rabbits from two independent experiments (three rabbits each) performed on different days. Individual values and geometric means of bacteria recovered from vegetations are shown. Identical symbols indicate bacteria recovered from the same animal. Log-transformed values were analyzed by paired Student’s t-test. *, P ≤ 0.002.

To further investigate the role of AsaA in binding to other host components present on damaged cardiac surfaces [42, 43], we tested the ability of an IE12 *asaA* mutant to bind to immobilized human fibrinogen and fibronectin. While binding to fibrinogen was not significantly affected for the *asaA* mutant, binding to fibronectin was significantly increased (Fig 8). While this may be the result of more efficient exposure of additional adhesins upon removal of AsaA, these results demonstrate that AsaA does not play a role in binding to fibrinogen or fibronectin.

**Fig 8.**
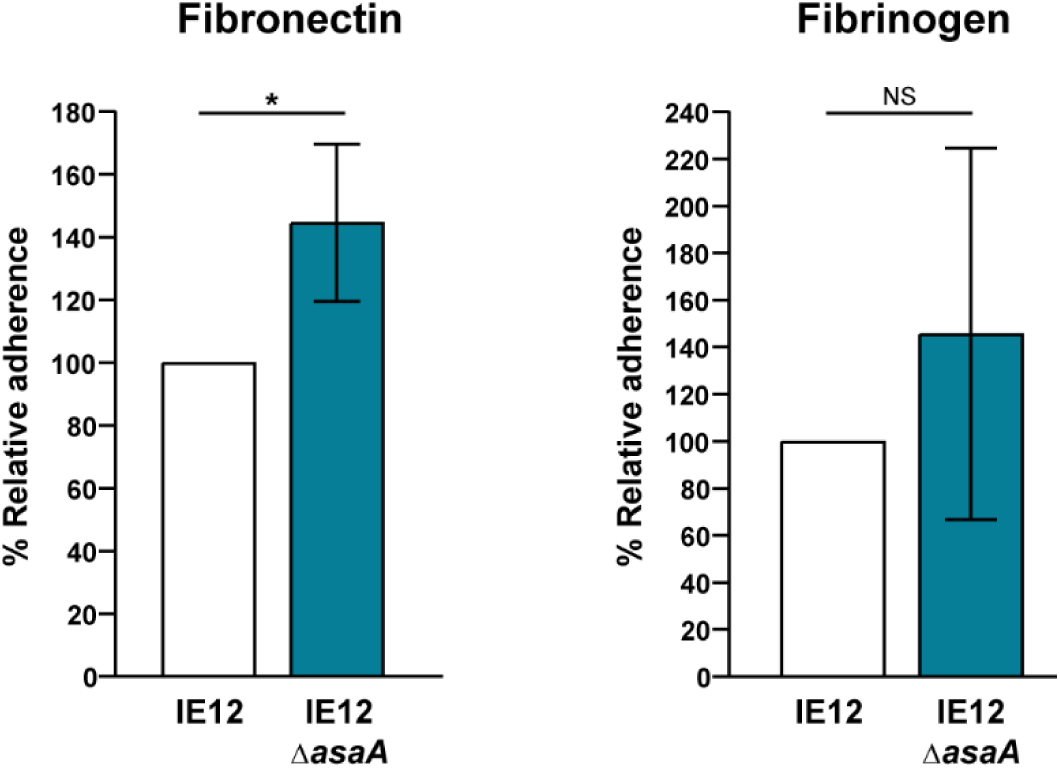
AsaA is not required for binding to fibrinogen or fibronectin. The IE12 *asaA* mutant displayed increased binding to immobilized fibronectin, while binding to fibrinogen was not significantly different from that of IE12. Adherence is expressed as a percentage relative to binding of IE12. Values are the means for at least three independent experiments, each performed in triplicate, ± SD. Statistical significance was tested by two-tailed Student’s t-test. *, P ≤ 0.02; NS, not significant.

### AsaA is present in other IE-causing bacterial species

Most studies of bacterial-platelet interactions have focused on a limited number of bacterial species, such as *S. gordonii* and *S. aureus* [44]. However, there are other bacterial species that cause IE through unknown mechanisms. To investigate if AsaA orthologues were present in other IE-causing species, we performed BLAST searches using the full-length amino acid sequence of *S. oralis* subsp. *oralis* AsaA as a query. AsaA orthologues were identified in other IE-causing bacterial species including *S. mitis, Staphylococcus pasteuri, Granulicatella elegans* and *G. haemolysans* [45-55]. Overall, the protein architecture of these AsaA orthologues is similar to that of *S. oralis* AsaA. They all contain a secretion signal, an NRR, DUF1542 repeats and a cell-wall anchoring motif. The NRR of *S. oralis* AsaA consists of a predicted FIVAR/CBM domain followed by two Siglec-like and Unique domains. The same domains were predicted in the AsaA orthologues in *G. haemolysans* M341, *S. pasteuri* 915_SPAS and *G. elegans* ATCC700633, although in the latter the N-terminal region of the NRR only shares structural homology with the FIVAR domain. Notably, we identified three Siglec-like and Unique domains in the NRR of the AsaA orthologue in *S. mitis* B6 (Fig 9A).

**Fig 9.**
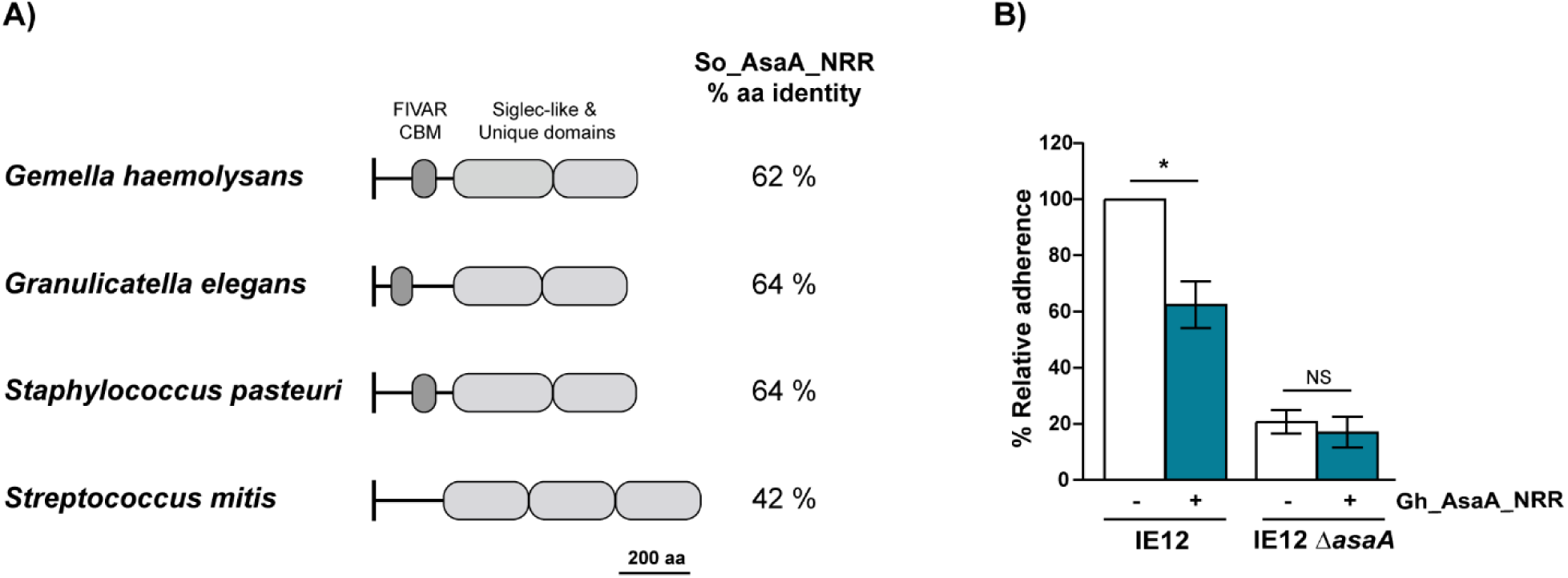
AsaA can serve as an adhesin in other IE-causing bacterial species. (A) Schematic illustrating the predicted protein architecture of the NRR of AsaA orthologues from *G. haemolysans* M341 (EGF87895.1), *G. elegans* ATCC700633 (EEW93886.2), *S. pasteuri* 915_SPAS (WP_048803571.1) and *S. mitis* B6 (CBJ22549.1). The percentage of amino acid identity to the NRR of AsaA from *S. oralis* subsp. *oralis* IE12 is shown to the right of the schematic. (B) Adhesion to platelets of IE12 and the *asaA* mutant in the presence of 5 µM Gh_AsaA_NRR. Adherence is expressed as a percentage relative to binding of the same strain in the absence of Gh_AsaA_NRR. Values are the means for at least three independent experiments, each performed in triplicate, ± SD. Statistical significance was tested by two-tailed Student’s t-test. *, P ≤ 0.0015; NS, not significant.

As a proof of principle that AsaA orthologues can also act as adhesins, we demonstrated that the NRR of AsaA from *G. haemolysans* M341 (Gh_AsaA_NRR), significantly reduced binding of IE12 to platelets. This reduction in binding was AsaA-dependent as adherence of the IE12 *asaA* mutant was not further reduced by the addition of Gh_AsaA_NRR (Fig 9B). These results demonstrate that AsaA from *G. haemolysans* can serve as a sialic acid-binding adhesin. Thus, this novel mechanism is potentially relevant to the pathogenesis of multiple IE-causing bacterial species.

## Discussion

Adhesion to host surfaces is a key step in bacterial pathogenesis. As such, bacterial binding to platelets plays an important role in the development of IE [8]. Initial studies in streptococci demonstrated that these interactions are mediated by SRRPs binding to sialic acid on platelets [9-15]. However, we previously demonstrate that the *S. oralis* subsp. *oralis* SRRP, Fap1, also mediates binding to the cryptic receptor β-1,4-linked galactose. This work revealed that these interactions are more complex than previously appreciated. Here, we confirm sialic acid and β-1,4-linked galactose as platelet receptors for *S. oralis* subsp. *oralis*. Nevertheless, our data demonstrate that SRRPs are not essential for binding these carbohydrates. Instead, we identified a novel sialic acid-binding adhesin, here named AsaA, which contributes to *S. oralis* subsp. *oralis* colonization of vegetations in an animal model of IE. The identification of AsaA-orthologues in additional IE-causing species and the demonstration that one of these proteins can competitively inhibit adhesion of a sialic acid-binding strain suggest that this novel adhesin contributes to the pathogenesis of multiple bacterial species that cause IE.

Previous reports have shown that binding of streptococci to platelets glycans is mediated by SRRPs [9-15]. Consistent with this, a previous study identified SRRPs in all 19 *S. gordonii* and *S. sanguinis* genomes analyzed [56]. In contrast, our results demonstrated that SRRPs are not present in all *S. oralis* subsp. *oralis* IE-isolates screened. While the requirement of SRRPs to bind platelet glycans may be specific for certain streptococcal species, our results stablish that SRRPs are not essential for binding of *S. oralis* subsp. *oralis* to sialic acid and β-1,4-linked galactose.

*S. oralis* subsp. *oralis* IE-isolates lacking SRRPs showed different binding dynamics to β-1,4-linked galactose than isolates encoding Fap1. While binding of Fap1-encoding isolates is reduced upon neuraminidase treatment (Fig2B and [9]), that of the SRRP-negative isolates IE1 and IE18 was either unaffected or increased, respectively (Fig 2A). This suggests that isolates lacking SRRPs bind to carbohydrates underlying sialic acid with higher affinity than those dependent on Fap1. This novel binding mechanism requires a SrtA-associated protein. Through comparative genomics and gene distribution analysis, we identified a Csh-like protein as a putative *S. oralis* β-1,4-linked galactose-binding protein.

Csh-like proteins are a widespread family of fibril-forming surface proteins found in oral streptococci such as *S. gordonii, S. oralis, S. mitis and S. sanguinis*. These proteins consist of an antigenically variable NRR and a conserved repetitive domain [57, 58]. The Csh-like protein identified in IE18 shares the highest amino acid identity (76%) with CshA from *S. gordonii*, which mediates lactose sensitive interspecies interactions and possesses a binding domain with a lectin-like fold (NR2) [33, 35, 36, 59]. The presence of Csh-like correlated with binding of SRRP-negative IE-isolates to β-1,4-linked galactose; however, we could not confirm a role for Csh-like in binding this carbohydrate. A *csh-*like mutant displayed a marked aggregation phenotype that interfered with binding assays; the reason for this aggregation phenotype is unknown but could be due to changes in cell hydrophobicity or to the exposure of other surface adhesins involved in aggregation. As an alternative approach, we tested the ability of the recombinantly expressed NRR of Csh-like to competitively inhibit binding to β-1,4-linked galactose; however, despite being properly folded, it did not reduce binding significantly. While our data do not support the hypothesis that Csh-like mediates binding of *S. oralis* to β-1,4-linked galactose, we cannot rule out this possibility.

In this work, we identified a novel sialic acid-binding adhesin, named AsaA, that is present in *S. oralis* subsp. *oralis* IE-isolates lacking SRRPs. An *asaA* mutant was reduced in binding to platelets, indicating that AsaA is required for adhesion. Additionally, we established that binding was sialic acid-dependent as removal of this carbohydrate did not further reduce binding of the mutant. We demonstrated that binding to platelets requires the AsaA Siglec-like and Unique domains, as a deletion mutant lacking these domains was reduced in adhesion. Finally, we established the role of AsaA as an adhesin by demonstrating that the recombinantly expressed NRR, specifically the Siglec-like and Unique domains, competitively inhibited adhesion of *S. oralis* subsp. *oralis* and directly bound to sialic acid on platelets.

In the most extensively studied SRRPs, Fap1, GspB, Hsa and SrpA, sialic acid interactions are mediated by a single Siglec-like domain [9-11, 15, 19-21]. This domain is invariably followed by a Unique domain, proposed to allosterically modulate the conformation of the Siglec-like domain [21]. In these SRRPs, Siglec-like domains have a semi-conserved YTRY motif containing an arginine residue essential for sialic acid binding [9-11, 15, 21, 22]. AsaA contains two Siglec-like and two Unique domains. AsaA_Siglec1_ contains a non-canonical YTRY motif (GTRY), which includes the arginine required for sialic acid binding. However, AsaA_Siglec2_ does not possess a YTRY motif. Two previously characterized SRRPs, FapC and SK1, also possess two Siglec-like and two Unique domains. In FapC, only the first Siglec-like domain has the conserved arginine; however, it is not essential for FapC binding to sialic acid on salivary proteins [39]. In SK1, neither of the Siglec-like domains has a conserved arginine and yet the protein binds sialic acid. How Siglec-like domains lacking the arginine of the YTRY motif bind sialic acid is unknown; however, a recent study suggests that additional interactions can occur between Siglec-like domains and sialoglycans through three flexible loops [20]. Together, these data suggest that both AsaA Siglec-like domains bind sialic acid. The impact of having two tandem repeats of Siglec-like and Unique domains is also unknown. However, previous studies showed that increasing the number of binding domains can increase affinity [29, 60]. Additionally, the NRR of SK1, containing two Siglec-like and Unique domains, binds a wider variety of sialoglycans than characterized SRRPs containing only one Siglec-like and Unique domain [21]. Hence, it is possible that increasing the number of Siglec-like and Unique domains broadens protein selectivity and impacts tropism, by expanding the range of host receptors than can be bound.

As indicated by their name, FIVAR domains are found in various bacterial proteins. FIVAR domains have been found in some CAZymes (Carbohydrate-active enzymes), and hence they are referred to as putative, yet uncharacterized, CBMs [61-65]. Since these domains share structural homology with heparin binding modules, it was suggested that they could bind negatively charged polysaccharides [63]. In agreement with the hypothesis that FIVAR domains bind carbohydrates, this region of AsaA was predicted to share structural homology with a putative CBM present in an endo-β-*N*-acetylglucosaminidase (EndoS) from *Streptococcus pyogenes* (PDB: 4NUZ) [66]. The putative CBM from EndoS shares structural homology to CBM62, which binds galactose-containing polysaccharides [66]. Although the ligands of the EndoS CBM remain unknown, nuclear magnetic resonance analysis showed potential low affinity binding to D-galactose [67]. These data suggest that, like Fap1, AsaA is involved in binding β-1,4-linked galactose exposed by neuraminidase cleavage of sialic acid [9]. However, an *asaA* mutant did not reduce binding of IE18 to neuraminidase pretreated platelets (S9 Fig). Additionally, the fact that neuraminidase pretreated platelets, which have exposed β-1,4-linked galactose, did not bind to the immobilized AsaA_NRR rules out a role for AsaA in binding to β-1,4-linked galactose. Although the role of the FIVAR domain is unclear, in our deletion mutants (Fig 5D) and platelet binding assay (Fig 6D) the presence of this domain tends to increase platelet binding. These data did not reach significance (P ≤ 0.250 and 0.09, respectively), therefore further studies are required to define the role of the FIVAR domain in AsaA.

FIVAR domains are also found in adhesive molecules such as Ebh from *S. aureus* and Embp from *S. epidermidis* [38, 68-70], which possess more than 40 FIVAR domains. Tandem FIVAR domains directly interact with the glycoprotein fibronectin [38]. However, we demonstrated that AsaA, which has only one FIVAR domain, is not involved in binding to fibronectin; whether this activity requires the presence of several FIVAR domains remains to be determined.

According to Lin *et al*, DUF1542 is the most repeated domain in streptococci [71]. Although Ebh and Embp consist mainly of FIVAR domains, these adhesins also possess four and seven DUF1542 domains, respectively [38, 68]. Additionally, *S. aureus* SasC, *Lactobacillus rhamnosus* MabA and *S. pyogenes* Epf possess 17, 26 and 18 DUF1542 domains, respectively [40, 72, 73]. Previous studies demonstrated that the DUF1542 domains present in SasC and Epf are not involved in adhesion, cell aggregation or biofilm enhancement [40, 72]. Instead, these domains form long, flexible, thin, fiber-like structures when overexpressed [40]. Thus, it is likely that AsaA forms a surface fibril.

A protein architecture similar to that of AsaA, consisting of an NRR followed by a DUF1542 domain stalk, is also found in Epf, MabA and SasC [40, 72, 73]. However, as observed in the SRRP-family, each adhesin possesses a unique NRR. The NRR of Epf is comprised of two subdomains, one is structurally similar to CBMs and the other adopts a fibronectin type III fold [40]. The structure of the NRRs of MabA and SasC are unknown; however, HHPred searches identified structural similarity between the NRR of MabA and the CBM-like domain of Epf [37, 40]. We propose that AsaA, Epf, SasC and MabA form part of a novel family of bacterial adhesins, here named DRAs (DUF1542-Repeat Adhesins), with a similar protein architecture.

Different NRRs are present in both DRAs and SRRPs, suggesting that these protein families can be a source of intra and interspecies diversity in receptor binding. Horizontal gene transfer between these organisms could lead to replacement of binding domains. The repeat domains in each protein family (Serine-rich and DUF1542) are not involved in adhesion, but are proposed to function as a stalk that helps the adhesive NRR protrude beyond the cell surface [40, 74, 75]. However, these domains are clearly different and may have a distinct impact on the biology of the organism. Serine-rich repeats are heavily glycosylated, which is important in biofilm development [76]. Additionally, these glycosylated adhesins require a specialized accessory Sec-System for their export [17]. No accessory Sec-system was found in the AsaA-encoding isolates, additionally, AsaA possess a YSIRK signal peptide that may drive its export through the general Sec-system. These data suggest that AsaA is not glycosylated and its synthesis and export is likely less energetically expensive than that of SRRPs. The expression of SRRPs has also been associated with high cell hydrophobicity [77]; therefore, SRR and DUF1542 domains will likely impact the biology of the organism in ways that are yet to be appreciated. Further experiments, beyond the scope of this study, are underway to establish the role of the repeat domains and their biological implications.

Adhesion is a key bacterial process during to host colonization, and hence it is linked to the virulence potential of bacteria. In streptococci, the expression of SRRPs is important for virulence in animal models of IE [15, 23-25, 78]. This phenotype is attributed to the ability of SRRPs to bind sialic acid on platelets [15]. Likewise, our data indicate that *S. oralis* AsaA is important for the colonization of vegetations in an *in vivo* model of IE. We observed a 3-fold reduction in the number of mutant bacteria recovered from the vegetations. Although a direct comparison with other *in vivo* studies is impractical due to variations in the models employed, this result is consistent with previous observations [14, 25, 78]. Additionally, the reduction in *asaA* mutant bacteria recovered is likely associated with the inability of the *asaA* mutant to bind sialic acid on platelets. In addition to platelets, other host factors such as fibrinogen and fibronectin are present on the damaged valve surface and contribute to initial bacterial binding [42, 43]. An *asaA* mutant was increased in binding to fibronectin, likely due to the absence of AsaA exposing other adhesins. This increased binding to other host factors may have partially compensated for the loss of AsaA *in vivo*. Our data demonstrate that AsaA contributes to the colonization of vegetations; however, more studies will be required to determine the impact on the course of disease.

We identified AsaA orthologues in other IE-causing species including *G. haemolysans, G. elegans, S. pasteuri* and *S. mitis* [45, 47, 48, 52, 53]. A previous study identified DUF1542-containing proteins in five *S. mitis* strains [79]. Although these were not characterized, we used Pfam and HHpred to determine that these proteins are AsaA orthologues, possessing Siglec-like and Unique domains in the N-terminal region. The overall architecture of these proteins is similar, however, some display differences in the NRR domains and the number of DUF1542 repeats. The AsaA orthologue in *S. mitis* B6 lacks the FIVAR/CBM domain, suggesting it is dispensable for protein function. Of note, the *S. mitis* AsaA orthologue contains three distinct Siglec-like and Unique domains in the NRR; to our knowledge, this is the first report of a streptococcal adhesin predicted to contain three such domains. These differences will likely impact the binding affinity and selectivity as discussed above.

The ability of the *G. haemolysans* AsaA orthologue (Gh_AsaA) to act as an adhesin was demonstrated by the reduction of *S. oralis* adhesion in the presence of the recombinantly expressed Gh_AsaA_NRR. The 40% reduction in binding to platelets was less than that observed in the presence of So_AsaA_NRR, the recombinantly expressed NRR of *S. oralis* AsaA. This result could be explained by variation in binding specificity or affinity derived from differences in the amino acid sequence. These results suggest the novel sialic acid-binding adhesin AsaA is a virulence factor important for the pathogenesis of multiple bacterial species that cause IE.

In summary, this study identified a novel sialic acid-binding adhesin that contributes to the virulence of *S. oralis* subsp. *oralis*. AsaA is broadly distributed and likely relevant to the pathogenesis of multiple IE-causing species. We propose that AsaA forms part of a novel family of bacterial adhesins that employ DUF1542 repeats to extend different NRRs beyond the cell surface. The NRRs, which determine the receptors bound, are shared with at least some of those found in SRRPs. However, since the repeat regions of these two families of bacterial adhesins are different, we hypothesize that they will differentially impact traits such as hydrophobicity and biofilm formation. Finally, our work demonstrates that sialic acid is a more broadly conserved receptor than previously appreciated, highlighting the importance of this bacterial adhesion mechanism in the biology of these organisms.

## Materials and methods

### Bacterial strains and culture media

All bacterial strains and plasmids used in this study are listed in Table 1. The five *S. oralis* strains included in the study, confirmed as subsp. *oralis* by MLSA, were selected from a group of eight isolates for their ability to bind efficiently and reproducibly to platelets in a sialic-acid dependent manner. *S. oralis* subsp. *oralis* was grown on tryptic soy agar plates (TSA) supplemented with 5% sheep blood (Becton, Dickinson and Co.) at 37°C in a 5% CO_2_ atmosphere for 16 h. Liquid cultures were grown statically at 37°C in Todd-Hewitt broth (Becton, Dickinson and Co.) supplemented with 0.2% (w/v) yeast extract (Becton, Dickinson and Co.) (THY). *Escherichia coli* strains were grown aerobically in lysogeny broth (LB) at 37°C with constant shaking at 250 rpm. When necessary, *S. oralis* subsp. *oralis* growth media was supplemented with the appropriate antibiotics at the following concentrations: 200 µg/mL spectinomycin, 500 µg/mL kanamycin or 2.5 µg/mL chloramphenicol. *E. coli* growth media was supplemented with 100 µg/mL ampicillin, 50 µg/mL spectinomycin, 50 µg/mL kanamycin or 30 µg/mL chloramphenicol when required.

**Table 1.**
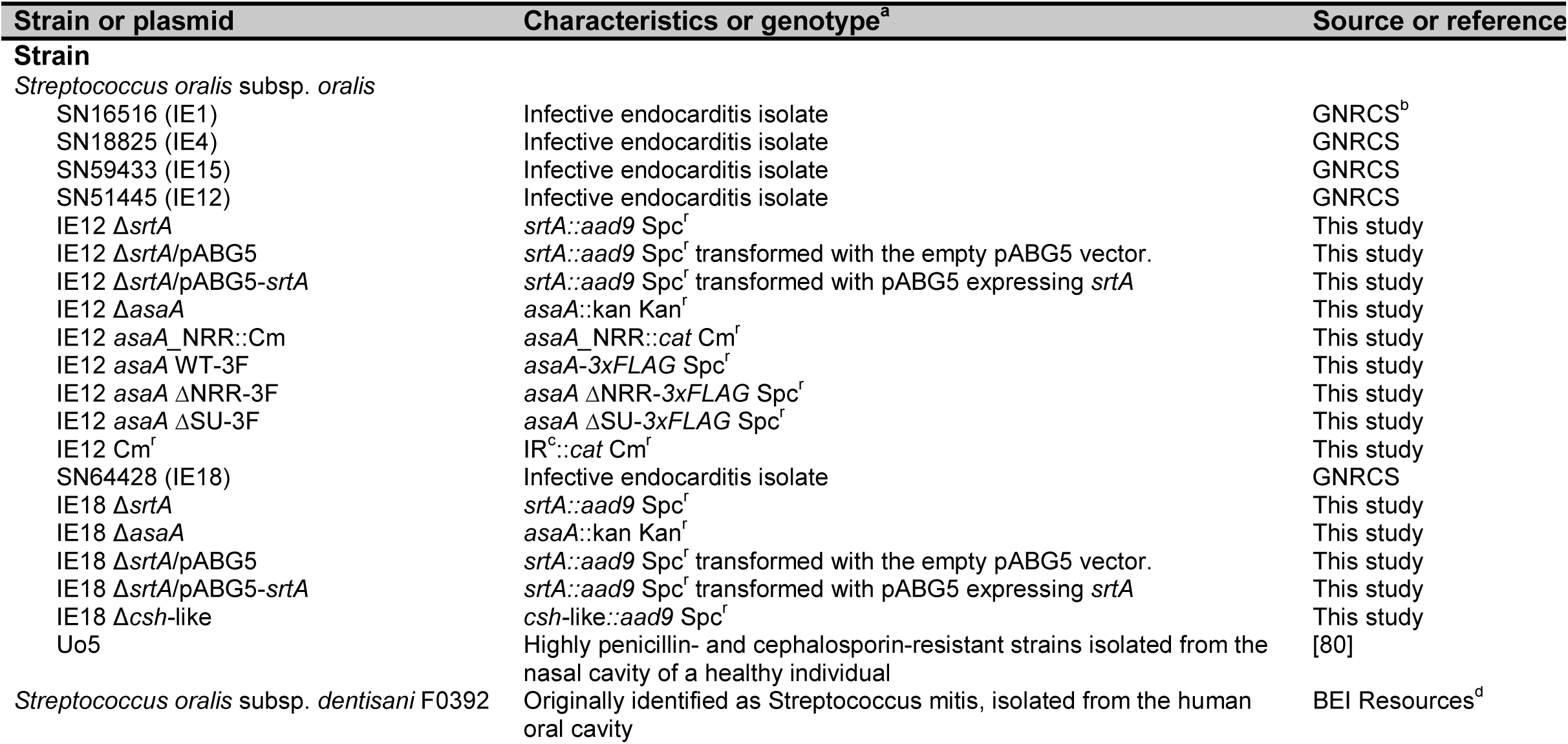
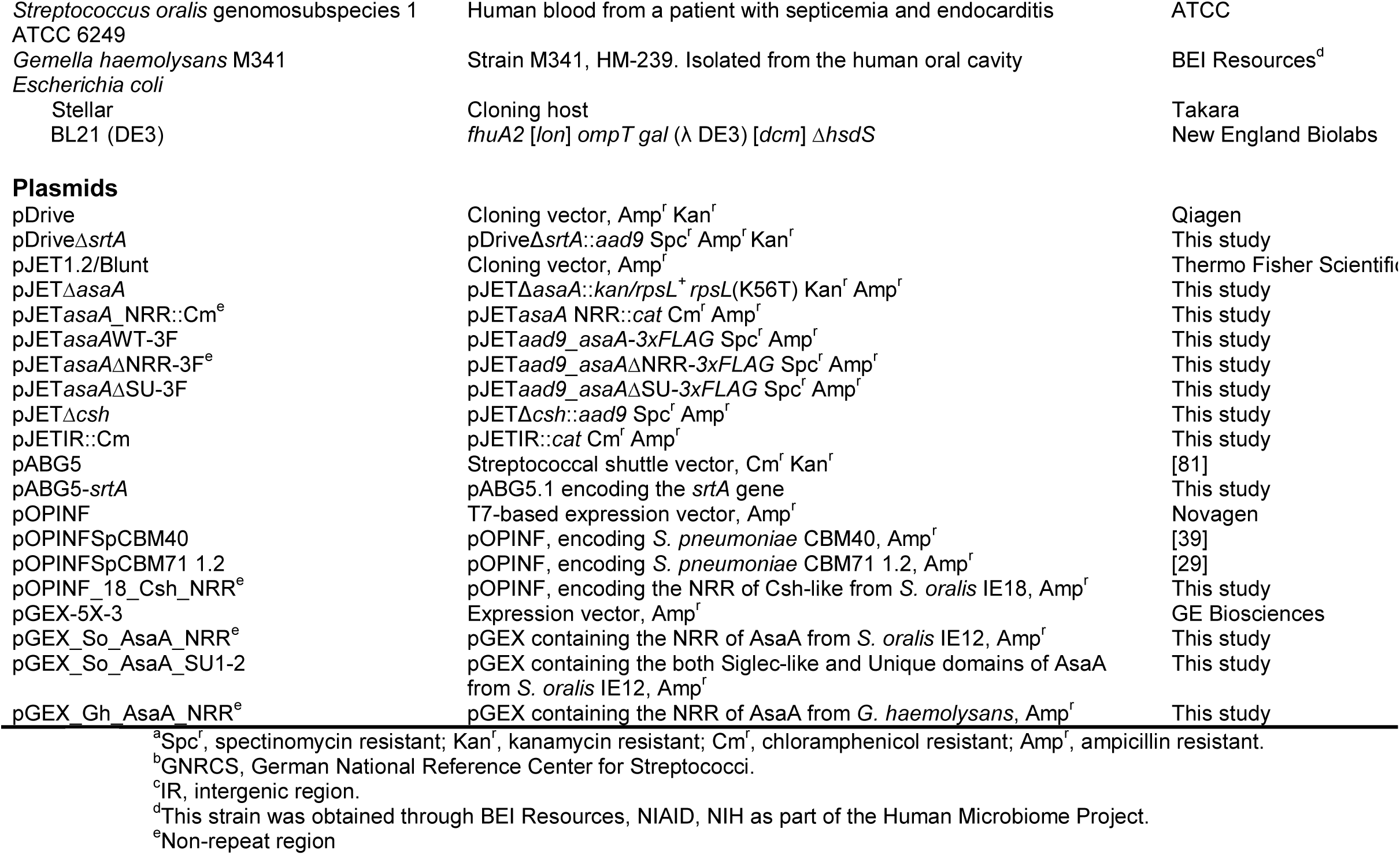
Strains and plasmids

For *in vivo* studies, bacteria were grown overnight at 37°C in brain heart infusion (BHI) broth under microaerobic conditions (6% O_2_, 7% H_2_, 7% CO_2_ and 80% N_2_) created with a programmable Anoxomat Mark II jar-filling system (AIG, Inc.); then diluted 10-fold into fresh, pre-warmed BHI and incubated statically at 37°C.

### Multilocus sequence analysis (MLSA)

*S. oralis* subsp. *oralis* isolates were identified using a previously published MLSA scheme [82]. Concatenated sequences of seven housekeeping genes were added to M. Kilian’s database of concatenated sequences from mitis group isolates. The species was confirmed and subspecies assigned using MEGA (version 6.06) software as previously described [83, 84].

### Genomic DNA isolation and sequencing

Genomic DNA (gDNA) was isolated as previously described [85] with slight modifications. Briefly, cells were grown in THY to an optical density at 600 nm (OD_600_) of 0.6 and harvested by centrifugation. The cell pellet was incubated at 37°C with 150 U of mutanolysin (Sigma) followed by treatment with RNAse A/N1 (100 µg/mL, Thermo Scientific), Proteinase K (100 µg/mL) and N-lauryl sarcosine (final concentration 1.5%). Two Phenol:Chloroform:Isoamyl alcohol (25:24:1) extractions were performed followed by two extractions with Chloroform. DNA was precipitated with isopropanol and washed with cold 70% ethanol. Finally, samples that were sequenced by PacBio were processed in accordance with the Guidelines for Using a Salt:Chloroform Wash to Clean Up gDNA (Pacific Biosciences).

gDNA from *S. oralis* subsp. *oralis* strains IE12 and IE18 was sheared to 10 kb with the Diagenode Megaruptor 2 converted into a SMRTbell library (Pacific Biosciences) using the SMRTbell express template preparation standard protocol. A total of 2-3.5 µg of the recovered library was concentrated with 0.6X AMPure PB carboxylated paramagnetic beads (Pacific Biosciences). The libraries were prepared for sequencing with Sequel binding kit 3.0 (Pacific Biosciences) at a 3 pM final loading concentration. Each purified library complex was sequenced on one 1Mv2LR SMRT cell with a 6 h pre-extension and 20 h movie time using diffusion loading. Raw sequence data was processed in SMRT Link with the “Circular Consensus Sequencing” application to produce >Q20 consensus reads.

*S. oralis* subsp. *oralis* strain ATCC10557 was sequenced and assembled by the University of Washington PacBio Sequencing Services.

### Genome assembly and annotation

All >Q20 consensus reads from the IE12 and IE18 genomes were assembled using Canu v 1.8 assembler with the following parameters genomeSize=1.9m maxThreads=10 maxMemory=20g [86]. BBmap aligner from the BBTools package (https://jgi.doe.gov/data-and-tools/bbtools/) was used to calculate the general assembly statistics (contig length, coverage and GC content) by re-aligning all reads to the corresponding assembly. The assemblies resulted in a circular contig (2,113,732 bp, 407.25x coverage and 40.97% GC content) and two non-circular contigs (14,198 bp 745.15x coverage and 42.6% GC content; 2,092,871 bp, 726.19x coverage and 40.94% GC content) for the IE12 and IE18 genomes, respectively. For both genomes, completeness was estimated as 100% by the BUSCO 3.2.0 pipeline using the Firmicutes group orthologue database with translations of predicted open reading frames in each genome [87]. PROKKA version 1.12 was used for gene prediction and annotation, with SignalP for Gram-positive signal peptides prediction and Infernal for noncoding RNA annotation (--gram +/pos --rfam parameters, respectively) [88-90].

### Distribution of *fap1, secA2* and genes encoding potential carbohydrate binding adhesins

The presence of *fap1* and *secA2* in five *S. oralis* subsp. *oralis* IE-isolates was analyzed by Southern blotting using 2 µg of EcoRV digested gDNA as previously described [26]. Briefly, conserved internal fragments of *fap1* and three different *secA2* genes representing the different variants of this gene identified in *S. oralis*, were amplified using primers Sb.1 - Sb.2 and Sb.3 to Sb.8 (Table 2) and labeled with digoxigenin (DIG) using the PCR DIG probe synthesis kit (Roche Diagnostics). Membranes were hybridized with these probes at 65°C and then washed at 68°C with decreasing concentrations (from 2X to 0.2X) of SSC (150 mM NaCl, 15 mM sodium citrate). Bound probes were detected using the DIG nucleic acid detection kit. As a positive control, all membranes were re-probed with *secA*.

**Table 2.**
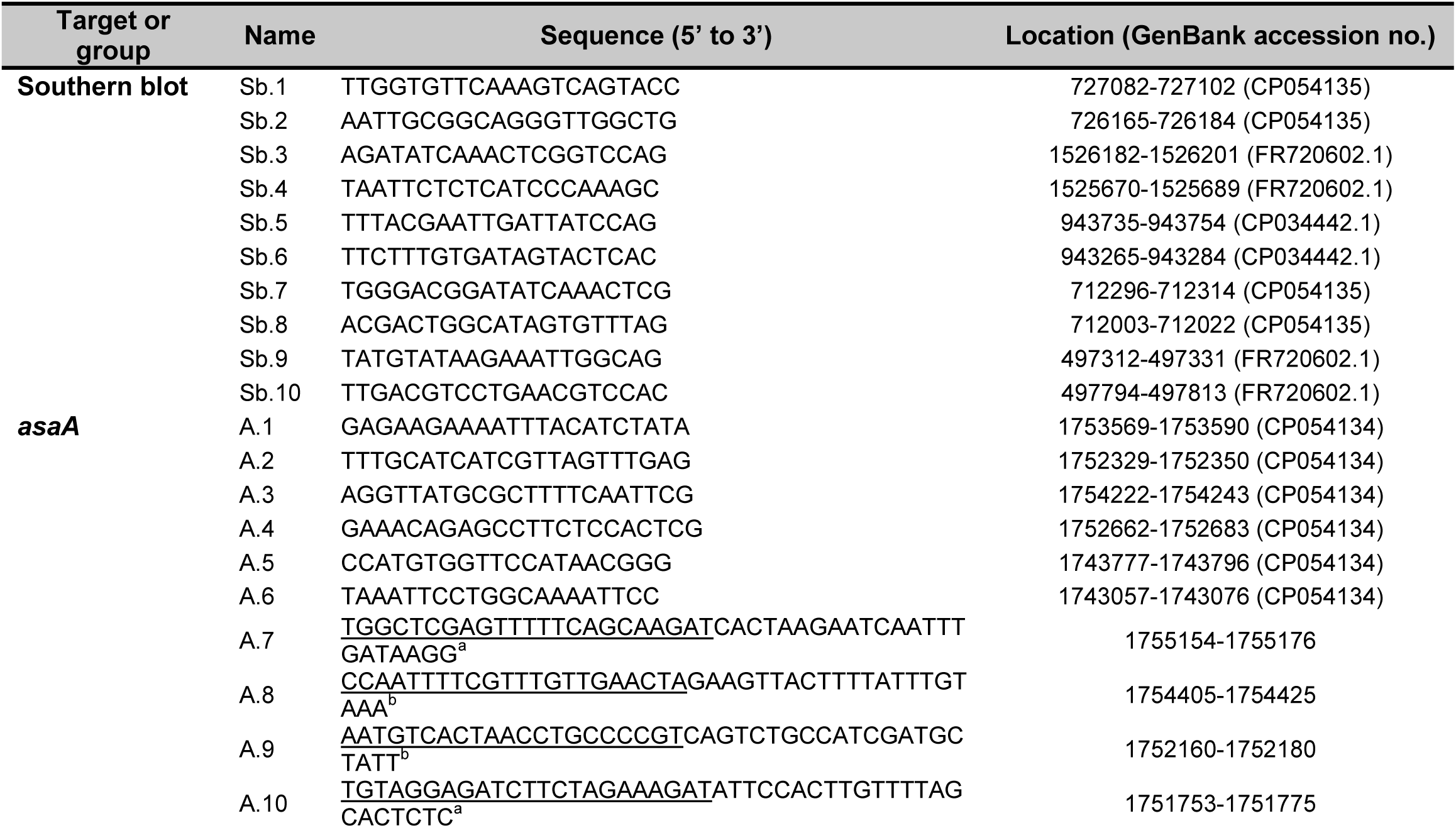
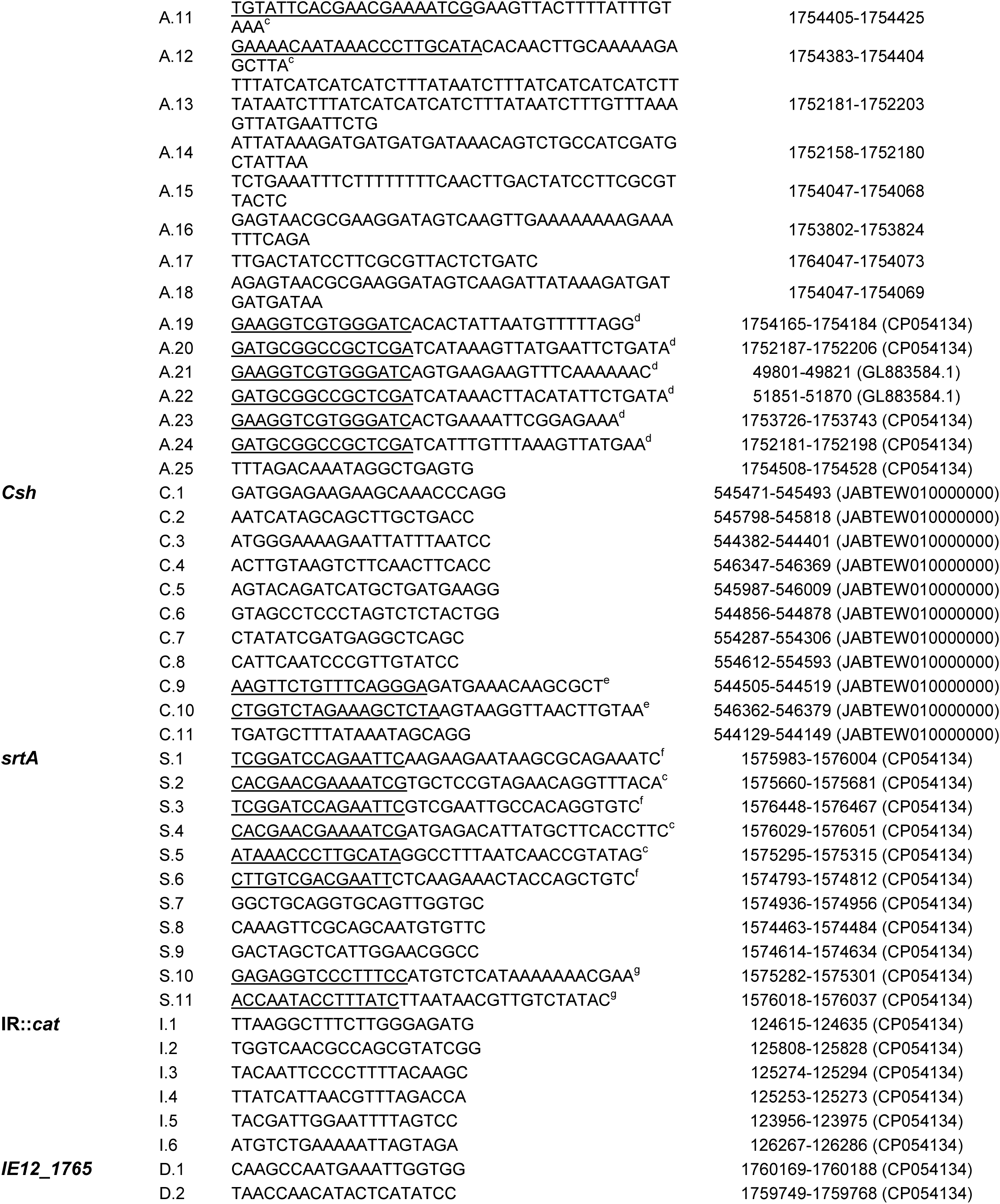
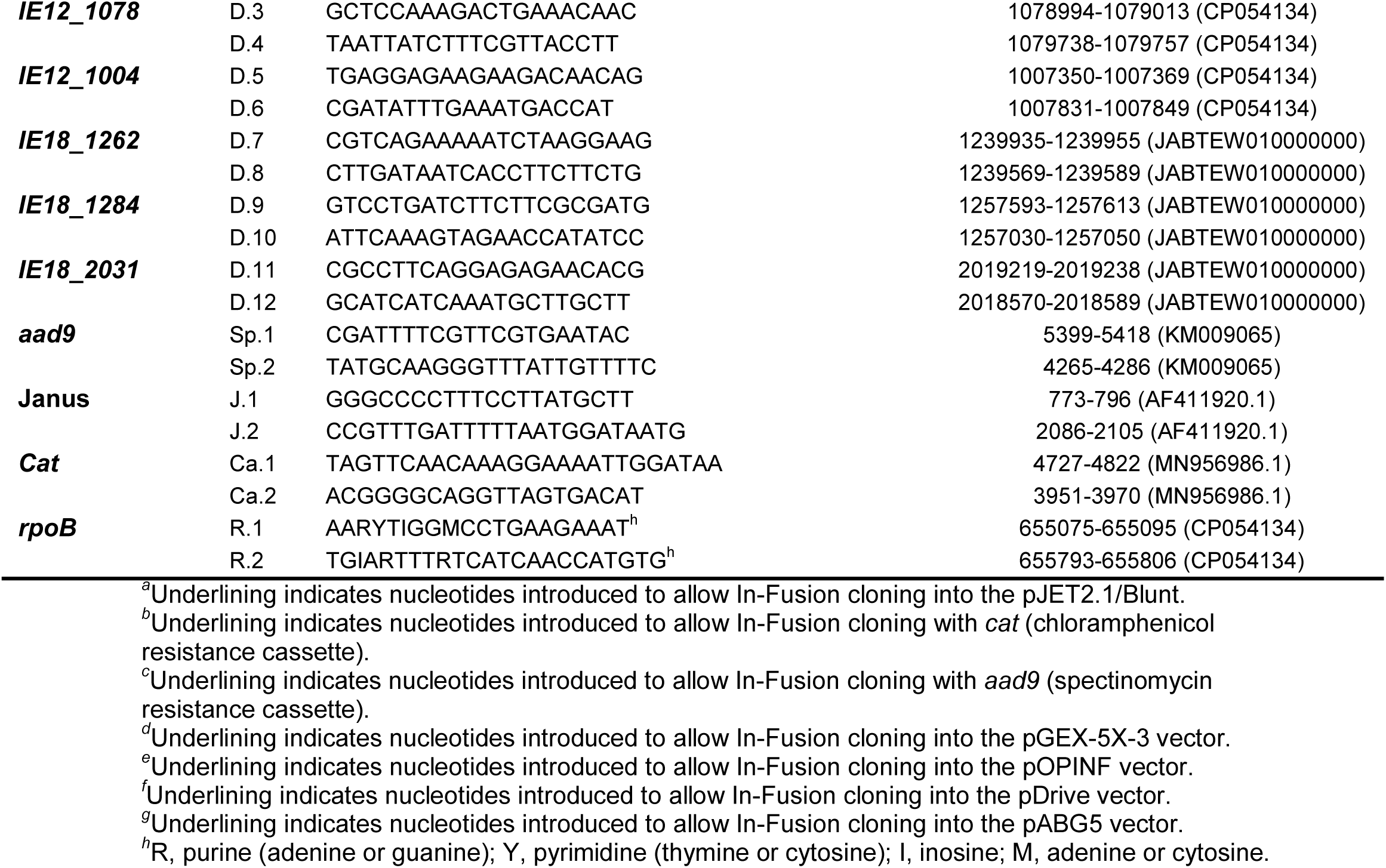
Primers used in this study

The presence of the genes encoding potential carbohydrate binding adhesins in the five different IE-isolates was determined by PCR using the corresponding primer pairs (Table 2).

### RNA isolation and RT-PCR

RNA isolation was conducted with the RNeasy mini kit (Qiagen) as previously reported [39]. Briefly, the cell pellet of 1 mL of bacterial culture grown to an OD_600_ of 0.4 was washed twice with phosphate-buffered saline (PBS) and resuspended in 540 µL of acid phenol containing 6 µL of 10% SDS. After flash-freezing, the samples were thawed at 70°C and then 540 µL of chloroform and 100 µL of Tris-EDTA buffer were added.

Samples were further incubated for 30 minutes at 70°C and vortexed every 5 minutes. RLT buffer was added to the sample, vortexed and centrifuged at 12,000 x g for 12 minutes. The aqueous phase was mixed with 70% ethanol, loaded into a RNeasy minicolumn (Qiagen) and further processed following the manufacturer’s instructions. RNA was treated with DNase I (Invitrogen) and further purified with the RNeasy mini kit. cDNA was synthesized from 1 µg of RNA using SuperScript II reverse transcriptase (Invitrogen) according to the manufacturer’s instructions.

Expression of *srtA, asaA* and *csh-like* was confirmed with the primer pairs S.1 - S.2, A.1 - A.2 and C.1 - C.2, respectively. To exclude possible polar effects caused by the deletion of *srtA, asaA* or *csh-like*, expression of the genes encoded downstream was confirmed using primer pairs S.7 - S.8, A.5 - A.6 and C.7 - C.8, respectively. In all cases, *rpoB* was used as a positive control (primers R.1 and R.2).

### Plasmid construction

The plasmid constructs used to generate the *S. oralis* subsp. *oralis* Δ*srtA*, Δ*asaA* and Δ*csh-like* mutants are described below. The regions upstream and downstream of the *srtA* gene were amplified from *S. oralis* subsp. *oralis* gDNA using the primer pairs S.3 - S.4, and S.5 - S.6, respectively. The spectinomycin cassette was amplified with primers Sp.1 and Sp.2. These three fragments were cloned into EcoRI-digested pDrive (Qiagen) using the In-Fusion HD EcoDry cloning kit (Takara Bio), resulting in the plasmid pDriveΔ*srtA*. The construct to generate the *asaA* mutant was generated by amplifying a 1.5 kb fragment, corresponding to the NRR of *asaA*, with primers A.3 and A.4. The product was blunt-end ligated into the pJET1.2/Blunt PCR cloning vector (Thermo Fisher Scientific). The resulting plasmid was digested with HincII, which removed ∼400 bp of *asaA*, then the Janus cassette was blunt-end ligated into this construct, generating the pJETΔ*asaA* plasmid. A 1.9 kb region, corresponding to the NRR-encoding region of *csh-like*, was amplified using primers C.3 and C.4, and blunt-end ligated into pJET1.2. The generated construct was used as a template for inverse PCR with primers C.5 and C.6. The resulting PCR product was blunt-end ligated to the spectinomycin resistance cassette, generating the pJETΔ*csh* plasmid. For complementation studies, the *srtA* gene was amplified by PCR using primers S.10 and S.11 and cloned by In-fusion in pABG5.

The plasmids used to generate the deletion mutants lacking the NRR or Siglec and Unique domains are described below. First, to generate a strain lacking the NRR (see below), we constructed the plasmid pJET*asaA_*NRR::Cm. To this end, two PCR products comprising ∼800 bp upstream of the promoter of *asaA* (Up) and ∼400 bp downstream of codon 704 (Dn) were amplified using primer pairs A.7 - A.8 and A.9 - A.10, respectively. The chloramphenicol cassette (Cm) was amplified using primers Ca.1 and Ca.2. These three PCR fragments (Up-Cm-Dn) were cloned into pJET1.2/Blunt using the In-Fusion HD EcoDry cloning kit. Next, we constructed a plasmid in which a spectinomycin resistance cassette was introduced before the promoter of *asaA*. We did so by cloning the following PCR products into pJET1.2/Blunt: a fragment comprising ∼800 bp upstream the *asaA* promoter, amplified using primers A.7 and A.11, the spectinomycin resistant cassette and a 2.7 kb DNA region (amplified using primers A.12 and A.10) comprising the promoter and the first 846 codons of *asaA*. The resulting construct was used as a template to amplify two PCR fragments using primer pairs A.7 - A.13 and A.14 - A.10 (primer A.13 and A.14 introduce a 3xFLAG tag) that were cloned by In-Fusion into pJET1.2/Blunt. The resulting plasmid was named pJET*asaA*WT-3F. Finally, to generate the pJET*asaA*ΔNRR-3F and pJET*asaA*ΔSU-3F constructs, two PCR fragments for each construct were amplified using gDNA from IE12 *asaA* WT-3F as a template with the following primer pairs: A.7 - A.15/A.16 - A.10 and A.7 - A.17/A.19 - A.10, respectively. These PCR fragments were cloned by In-Fusion into pJET1.2/Blunt.

For co-infection studies in a rabbit model of IE we generated an isogenic IE12 strain resistant to chloramphenicol (IE12 Cm^r^). To that end, we used primers I.1 and I.2 to amplify a conserved intergenic region (IR) between two convergent genes encoding: a TrkH family potassium uptake protein (TrkG) and an oligopeptide ABC transporter ATP-binding protein (OppF). The product was blunt-end ligated into pJET1.2 and used as a template for inverse PCR with primers I.3 and I.4. The resulting PCR product was blunt-end ligated to the chloramphenicol resistance cassette (pJETIR::Cm). For protein purification, pGEX_So_AsaA_NRR, pGEX_So_AsaA_SU_1-2 and pGEX_Gh_AsaA_NRR were constructed as follows. The full-length NRRs of AsaA from *S. oralis* subsp. *oralis* IE12 and *G. haemolysans* M341 (comprising amino acids 37 to 702 and 50 to 739, respectively), were amplified from gDNA using primers A.19 - A.20 and A.21 - A.22, respectively. A region comprising both Siglec-like and Unique domains of So_AsaA (amino acids 179-714) was amplified using primers A.23 - A.24. The resulting PCR products were cloned by In Fusion into pGEX-5X-3, which allows the expression of N-terminally GST-tagged proteins. The construct pOPINF_18_Csh_NRR was design to express the NRR of the Csh-like protein with an N-terminal His-tag. Briefly, a PCR fragment, comprising codons 42 to 666, was amplified using primers C.9 and C.10 and cloned into pOPINF using the In-Fusion HD EcoDry cloning kit (Takara Bio).

All constructs were confirmed by PCR and sequencing to verify that no spurious mutations had been introduced.

### Generation of mutants in *S. oralis* subsp. *oralis*

IE12 Δ*srtA*, IE12 Δ*asaA* and IE12 Cm^r^ were generated by transforming the plasmids pDriveΔ*srtA*, pJETΔ*asaA* and pJETIR::Cm into IE12. While for the IE18 *srtA* and IE18 *csh-like* mutants, pDriveΔ*srtA* and pJETΔ*csh* plasmids were transformed into IE18. Briefly, *S. oralis* subsp. *oralis* strains were grown in C+Y media to an OD_600_ of 0.1 and then diluted in fresh media (1:20). Competence was induced by the addition of a competence stimulating peptide (2 µg/mL) and 1 mM CaCl_2_ [9]. Approximately 100 ng of plasmid was added and the transformations were then incubated for 2 h at 37°C. Transformants were selected on TSA plates with the appropriate antibiotics. All desired mutations were confirmed by PCR using flanking primers (A.25, C.11, S.9 and I.5) in combination with primers for the antibiotic resistance cassette included in each mutant (J.1, Sp.2, Sp1 and Ca.2).

To complement the *srtA* mutants, the plasmid pABG5-*srtA* was transformed by electroporation as previously described [91]. Briefly, cells were grown in THY supplemented with 20 mM glycine to an OD_600_ of 0.2, harvested by centrifugation and resuspended in ice-cold electroporation medium (272 mM glucose, 1 mM MgCl_2_, pH 6.5). DNA was added to an aliquot of cells and transformed by electroporation. After 2 h incubation in THY at 37°C cells were plated into TSA plates with the appropriate antibiotics.

To generate the deletion mutants lacking the NRR or Siglec-like and Unique domains of AsaA we first generated an intermediate strain in which the NRR of AsaA was replaced with the chloramphenicol cassette. To this end, the pJET*asaA_*NRR::Cm construct was transformed into *S. oralis* subsp. *oralis* IE12 as described above. The resulting strain (IE12 *asaA_*NRR::Cm) was then transformed with the pJET*asaA*WT-3F, pJET*asaA*ΔNRR-3F or pJET*asaA*ΔSU-3F plasmids. The resulting colonies were selected for the resistance to spectinomycin and sensitivity to chloramphenicol. Desired mutations were confirmed by sequencing.

The genetic background of all mutants was confirmed by repetitive extragenic palindromic PCR [92]. Any possible growth defects were excluded by conducting growth assays on rich medium as previously described [39].

### Protein overproduction and purification

The N-terminally His-tagged CBM40 and the two tandem CBM71 domains (CBM71-1.2) from *Streptococcus pneumoniae* neuraminidase NanA and β-galactosidase BgaA, respectively, were purified by nickel affinity chromatography as described previously [29, 39].

To purify GST or the GST-tagged NRRs of So_AsaA and Gh_AsaA, plasmids pGEX-5X-3, pGEX_So_AsaA_NRR pGEX_So_AsaA_SU1-2 or pGEX_Gh_AsaA_NRR were transformed into the *E. coli* host strain BL21 (DE3) (Invitrogen). Overnight cultures of the transformed strains, grown at 37°C with constant shaking, were used to inoculate fresh LB containing the appropriate antibiotics. Growth was continued under the same conditions until an OD_600_ of 0.8 was reached. At this point, protein production was induced by the addition of isopropyl β-D-1-thiogalactopyranoside (IPTG) to a final concentration of 0.5 mM followed by a 4 h incubation at 30°C with constant shaking. Cells were harvested by centrifugation and lysed using a French Press. Cleared cell lysates were used for further purification by affinity chromatography. The cleared cell lysates were incubated overnight with glutathione-Sepharose 4B beads (GE Healthcare Life Sciences) in the presence of 5 mM dithiothreitol (DTT) at 4°C with gentle agitation. The beads were extensively washed with 50 mM Tris-HCl, 150 mM NaCl, pH 8.0. Finally, GST and GST-tagged proteins were eluted with 10 mM reduced glutathione.

The purification of the NRR of Csh-like was carried out as previously described for the NR2 of CshA [36]. Briefly, protein expression was induced by the addition of 1 mM IPTG for 16 h at 18°C. The His-tagged protein was purified from the cleared cell lysate by nickel affinity chromatography (Ni-NTA agarose beads, Invitrogen). After washing with increasing concentrations of imidazole, proteins were eluted with 200 mM imidazole in 20 mM Tris-HCl, 150 mM NaCl, pH 8.

Purified proteins were dialyzed overnight against PBS containing 1 mM DTT. Size and purity were verified by Coomassie blue-stained SDS-PAGE. Protein concentration was determined using the absorbance at 280 nm and the extinction coefficient.

Proper folding of the Csh_NRR was confirmed by CD-spectroscopy. CD-spectra of Csh_NRR in PBS was collected using a Jasco J-815 spectrometer, using a 1 mm path length cuvette, with a wavelength interval of 0.5 nm. Data was analyzed and plotted using CAPITO (CD Analysis and Plotting Tool) [93].

### Platelet isolation

Platelet isolation has been described previously and was performed with slight modifications [9]. Blood from healthy donors was collected in acid citrate dextrose solution (75 mM sodium citrate, 39 mM citric acid, 135 mM dextrose, pH 7.4). Pooled platelet rich plasma was collected after centrifugation of blood samples at 200 x g for 20 minutes and mixed 1:1 with HEP buffer (140 mM NaCl, 2.7 mM KCl, 3.8 mM HEPES, 5 mM EDTA, pH 7.4). Platelets were collected by centrifugation at 800 x g for 20 minutes with no brake applied and washed twice with platelet wash buffer (10 mM sodium citrate, 150 mM NaCl, 1 mM EDTA, 1% (w/v) dextrose, pH 7.4). Platelets were fixed with 1% paraformaldehyde for 10 minutes at room temperature and then washed three times with PBS. Platelets were resuspended in PBS at 1×10^7^ cells per mL. Flat-bottomed 96-well microtiter plates were coated with 100 µL of platelets for 1 h at 37°C. Control wells were coated with 1% (w/v) bovine serum albumin (BSA) in PBS. Unbound platelets were removed by two PBS washes. Platelets-coated plates were stored at 4°C until further use.

### Adherence assays

Where indicated, platelet-coated wells were pretreated with 0.005 U of *Clostridium perfringens* neuraminidase (Sigma) for 30 minutes at 37°C, control wells were incubated with PBS. After washing with PBS, all wells were blocked with 3% BSA in PBS for 1 h at 37°C.

Bacteria were grown in THY to an OD_600_ of 0.5 ± 0.05. Approximately 2×10^5^ bacteria in PBS were allowed to bind to fixed platelets at 37°C for 60 minutes. Three washes with PBS were performed to remove non-adherent bacteria and then adherent bacteria were lifted with 0.25% trypsin-1 mM EDTA at 37°C for 15 minutes. CFU counts were performed by serial dilution.

To test the effect of CBM40, CBM71-1.2, Csh_NRR, AsaA_NRR, AsaA_SU1-2 and Gh_AsaA_NRR on bacterial binding to platelets, soluble protein was added to the bacterial inoculum at the concentrations indicated in each experiment.

### Platelet binding to immobilized protein

Binding assays were performed in 96-well plates (Nunc MaxiSorp) coated overnight at 4°C with 100 µL of 5 µM AsaA_NRR, AsaA_SU_1-2 or GST. After washing to remove unbound protein with PBS, wells were blocked with 1% BSA at 37°C for 1 h. Two washes with PBS were performed before adding 100 µL of platelets (1×10^7^ platelets per mL) pretreated with PBS or neuraminidase. After 2 h of incubation at 37°C wells were washed three times with PBS and then fixed with 90% ethanol. Fixed platelets were stained with 0.1% crystal violet and imaged using a Nikon eclipse Ti inverted microscope with a 40x objective. The number of platelets in each field was quantified using ImageJ [94].

### Detection of AsaA-3xFLAG

The production of the AsaA deletion mutants was verified by western blot. All strains were grown to an OD_600_ of 0.5. After harvesting cells by centrifugation, the pellet was resuspended in PBS containing 1% sodium deoxycholate and 1% SDS and normalized according to the OD (PBS volume = OD_600_ x 400). Ten microliters of the whole cell lysates were spotted onto nitrocellulose membrane. After an hour, the membrane was blocked for 1 h with 1% BSA in PBS. Three washes with tTBS (50 mM Tris, 150 mM NaCl and 0.1% Tween 20) were performed before incubating the membrane with the anti-FLAG antibody 1:5000 for 1 h (Sigma). Next, after three washes with tTBS, the membrane was incubated with the secondary antibody 1:5000 for 1 h (HRP-coupled anti-rabbit IgG, Invitrogen). Protein detection was performed with the SuperSignal West Pico Plus kit according to the manufacturer’s instructions (Thermo Scientific).

Detection of the AsaA-3xFLAG deletion versions on the surface of IE12 was done with immunofluorescence. Cells were grown in THY until an OD_600_ of 0.5 was reached, then Nile red was added to the media (final concentration 1 ug/mL). After an additional 20 minute incubation, the cells were harvested by centrifugation. The cell pellets were washed three times with PBS and then fixed for 20 minutes with 40% methanol. Cells were washed with PBS and then incubated with anti-FLAG antibody 1:200 (Sigma) for 1 h at room temperature. After three washes with PBS cells were incubated with the secondary antibody 1:200 (Alexa Fluor 488, goat anti-rabbit IgG, Invitrogen) for 1 h. Finally, cells were washed three times and imaged on a LSM 800 confocal microscope (Zeiss, Germany). Images were rendered with Zeiss Zen software (Zeiss, Germany).

### Rabbit model of IE

The rabbit model employed for virulence assays has been described previously [41] and was performed with minor variations. Briefly, specific-pathogen-free, male New Zealand White rabbits weighing 2 - 3 kg and supplied by Charles Rivers Laboratories (experiment 1) or RSI Biotechnology (experiment 2) were used. Animals were anesthetized prior to surgery, which involved insertion of a catheter through the right carotid artery to the aortic valve or left ventricle to induce minor damage to the endothelium. Bacterial strains were cultured separately, then washed and combined in 0.5 mL PBS for co-inoculation of approximately 1×10^7^ CFU of each strain into a peripheral ear vein two days after surgery. Remaining cells were sonicated to disrupt clumps and chains and plated on BHI plates with selective antibiotics using an Eddy Jet 2 spiral plater (Neutec Group, Inc.). The following day, the animals were euthanized and vegetations in or near the aortic valves were removed, homogenized in PBS, sonicated, and plated, as for the inoculum. Plates were incubated at 37°C in Anoxomat jars for two days prior to counting colonies. Results were normalized to account for small (≤ 10%) differences in the inoculum sizes of the competing strains.

### Bacterial binding to immobilized fibrinogen and fibronectin

Assays were performed in 96-well plates (Nunc MaxiSorp) coated overnight at 4°C with 100 µL of fibrinogen (20 µM, Invitrogen) or fibronectin (50 µM, EMD Millipore Calbiochem); control wells were coated with 3% BSA. After washing to remove unbound protein, wells were blocked for 1 h at 37°C with 3% BSA. Approximately 2×10^5^ bacteria in PBS were allowed to bind at 37°C for 2 h. Wells were washed three times with PBS and then adherent bacteria were lifted with 0.25% trypsin-1 mM EDTA at 37°C for 15 minutes. CFU counts were performed by serial dilutions.

### *In silico* analysis

Protein homology and structure were predicted using the HHPred server [37]. The AsaA three-dimensional model was built using the SWISS-MODEL server [95]. Protein structures were edited for visualization using PyMOL (The PyMOL Molecular Graphics System, Version 2.0 Schrödinger, LLC). Multiple sequence alignments were carried out using Clustal Omega [96].

### Statistical analysis

*In vitro* data are presented as mean of at least three independent experiments ± standard deviation (SD). Data analysis was performed using GraphPad Prism. Two-tailed Student’s t-test were used to determine differences in bacterial and platelet adhesion. For rabbit studies, log-transformed values were analyzed by paired Student’s t-test.

### Accession number(s)

The MLSA sequences for the five *S. oralis* subsp. *oralis* isolates were submitted to GenBank and have the accession numbers MT551124 to MT551158. Genome assemblies of *S. oralis* subsp. oralis strains ATCC10557, IE12 and IE18 were deposited at GenBank under the accession numbers CP054135, CP054134 and JABTEW010000000, respectively. The data have been deposited with links to BioProject accession number PRJNA635656 at the NCBI BioProject database (https://www.ncbi.nlm.nih.gov/bioproject/).

### Ethics Statement

Blood draws involved the informed written consent of donors, with ethical approval obtained from the Institutional Review Board of Nationwide Children’s Hospital under protocol number IRB16-01173. For the endocarditis model, rabbits were premedicated with acepromazine, buprenorphine, and bupivacaine at the incision site, and then anesthetized for surgery with isoflurane and sevoflurane. Buprenorphine SR LAB (ZooPharm) was provided for post-surgical analgesia. Euthanasia was achieved by administration of Euthasol (pentobarbital sodium and phenytoin sodium solution; Virbac AH Inc.). All procedures performed in relation to the rabbit endocarditis model were approved by the Institutional Animal Care and Use Committee of Virginia Commonwealth University under protocol number AM10030, and were consistent with United States Department of Agriculture Animal Welfare Act and Regulations, the United States Office of Laboratory Animal Welfare Policies and Laws, and The Guide for the Care and Use of Laboratory Animals published by the National Research Council.

## Acknowledgments

We thank Shannon Green, Karina Kunka, Tanya Puccio, and Nicai Zollar for technical assistance with the rabbit studies. We are thankful to M. Kilian for providing an MLSA database to confirm species assignment. We thank Alicia Friedman of the Biophysical Interaction and Characterization Facility at The Ohio State University for her help with circular dichroism and Frank Robledo-Avila for his technical help with microscopy analysis. *Gemella haemolysans* M341 was obtained through BEI Resources, NIAID, NIH, as part of the Human Microbiome Project.

## Supporting information

**S1 Fig. *secA2* distribution parallels that of *fap1* in *S. oralis* subsp. *oralis* IE-isolates**. Southern blots of five different *S. oralis* subsp. *oralis* IE-isolates using three different DIG-labeled *secA2* probes amplified from *S. oralis* Uo5, F0392 and ATCC6249, each of which represents one of the three distinct *secA2* alleles identified within sequenced *S. oralis* strains. gDNA from the strains used to amplify each probe was used as a positive control for the appropriate blot (+).

**S2 Fig. Distribution of genes encoding potential carbohydrate binding adhesins**. The presence of the four genes shared between IE12 and IE18 (A) and those unique to IE18 (B) in all five *S. oralis* subsp. *oralis* IE-isolates was determined by PCR. The distribution of these genes was correlated with the ability of the IE-isolates to bind sialic acid and β-1,4-linked galactose. Amplification of *rpoB* was used as a positive control.

**S3 Fig. Expression of IE12_1764 and IE18_0557 is *S. oralis* subsp. *oralis***. Expression of IE12_1764 and IE18_0557 in IE12 and IE18, respectively, was analyzed by RT-PCR. In both cases, the expression of *rpoB* served as a positive control. To rule out DNA contamination, cDNA synthesis was performed in the absence of reverse transcriptase (-).

**S4 Fig. IE18 *csh-like* mutant forms aggregates**. Gram-staining of IE18 and IE18 Δ*csh-like* shows bacterial aggregates that could not be disrupted by vortexing. Representative images captured using a Nikon eclipse Ti inverted microscope with a 40x objective.

**S5 Fig. Csh_NRR is properly folded**. Representative CD spectra of the recombinantly expressed Csh_NRR at a concentration of 31.3 µM in PBS.

**S6 Fig. An independent IE12 *asaA* mutant is reduced in binding to platelets**. Adhesion of an independent IE12 *asaA* mutant to platelets pretreated with neuraminidase (N^PRE^) or PBS (-). Adherence is expressed as a percentage relative to binding of IE12 to untreated platelets. Values are the means for at least three independent experiments, each performed in triplicate, ± SD. Statistical significance was tested by two-tailed t Student’s t-test. *, P ≤ 0.0001. NS, not significant.

**S7 Fig. IE12 *asa*A WT-3F is reduce in binding to platelets**.

Adhesion to platelets of a strain expressing AsaA-3xFLAG (WT-3F) is reduced as compared to IE12. However, an *asaA* mutant is further reduced. Adherence is expressed as a percentage relative to binding of IE12. Values are the means for at least three independent experiments, each performed in triplicate, ± SD. Statistical significance was tested by two-tailed Student’s t-test. *, P ≤ 0.0007.

**S8 Fig. GST does not reduce binding of IE12 to platelets**. Unlike the addition of the recombinantly expressed So_AsaA_NRR (10 µM), the addition of 10 µM of GST alone did not significantly reduce binding of IE12 to platelets. Adherence is expressed as a percentage relative to binding of IE12 in the absence of So_AsaA_NRR or GST. Values are the means for at least three independent experiments, each performed in triplicate, ± SD. Statistical significance was tested by two-tailed Student’s t-test. *, P ≤ 0.0001; NS, not significant.

**S9 Fig. AsaA is not involved in IE18 binding to** β **1-4 linked galactose**. Adhesion of an IE18 *asaA* mutant to platelets pretreated with neuraminidase (N^PRE^). Adherence is expressed as a percentage relative to binding of IE18. Values are the means for at least three independent experiments, each performed in triplicate, ± SD. Statistical significance was tested by two-tailed t Student’s t-test. NS, not significant.

**S1 Table. Prediction of domains within the potential carbohydrate binding proteins**. The full amino acid sequence of the putative sialic-acid and β-1,4-linked galactose binding proteins was used to perform BLASTP and Pfam searches.

## References

1. Heller D, Helmerhorst EJ, Gower AC, Siqueira WL, Paster BJ, Oppenheim FG. Microbial Diversity in the Early In Vivo-Formed Dental Biofilm. Appl Environ Microbiol. 2016;82(6):1881–8.

2. Sulyanto RM, Thompson ZA, Beall CJ, Leys EJ, Griffen AL. The Predominant Oral Microbiota Is Acquired Early in an Organized Pattern. Sci Rep. 2019;9(1):10550.

3. Cahill TJ, Prendergast BD. Infective endocarditis. Lancet. 2016;387(10021):882–93.

4. Knox KW, Hunter N. The role of oral bacteria in the pathogenesis of infective endocarditis. Aust Dent J. 1991;36(4):286–92.

5. Lockhart PB, Brennan MT, Sasser HC, Fox PC, Paster BJ, Bahrani-Mougeot FK. Bacteremia associated with toothbrushing and dental extraction. Circulation. 2008;117(24):3118–25.

6. Holland TL, Baddour LM, Bayer AS, Hoen B, Miro JM, Fowler VG, Jr. Infective endocarditis. Nat Rev Dis Primers. 2016;2:16059.

7. Naveen Kumar V, van der Linden M, Menon T, Nitsche-Schmitz DP. Viridans and bovis group streptococci that cause infective endocarditis in two regions with contrasting epidemiology. Int J Med Microbiol. 2014;304(3-4):262–8.

8. Werdan K, Dietz S, Loffler B, Niemann S, Bushnaq H, Silber RE, et al. Mechanisms of infective endocarditis: pathogen-host interaction and risk states. Nat Rev Cardiol. 2014;11(1):35–50.

9. Singh AK, Woodiga SA, Grau MA, King SJ. Streptococcus oralis Neuraminidase Modulates Adherence to Multiple Carbohydrates on Platelets. Infect Immun. 2017;85(3).

10. Bensing BA, Loukachevitch LV, McCulloch KM, Yu H, Vann KR, Wawrzak Z, et al. Structural Basis for Sialoglycan Binding by the Streptococcus sanguinis SrpA Adhesin. J Biol Chem. 2016;291(14):7230–40.

11. Loukachevitch LV, Bensing BA, Yu H, Zeng J, Chen X, Sullam PM, et al. Structures of the Streptococcus sanguinis SrpA Binding Region with Human Sialoglycans Suggest Features of the Physiological Ligand. Biochemistry. 2016;55(42):5927–37.

12. Bensing BA, Lopez JA, Sullam PM. The Streptococcus gordonii surface proteins GspB and Hsa mediate binding to sialylated carbohydrate epitopes on the platelet membrane glycoprotein Ibalpha. Infect Immun. 2004;72(11):6528–37.

13. Takamatsu D, Bensing BA, Cheng H, Jarvis GA, Siboo IR, Lopez JA, et al. Binding of the Streptococcus gordonii surface glycoproteins GspB and Hsa to specific carbohydrate structures on platelet membrane glycoprotein Ibalpha. Mol Microbiol. 2005;58(2):380–92.

14. Siboo IR, Chambers HF, Sullam PM. Role of SraP, a Serine-Rich Surface Protein of Staphylococcus aureus, in binding to human platelets. Infect Immun. 2005;73(4):2273–80.

15. Pyburn TM, Bensing BA, Xiong YQ, Melancon BJ, Tomasiak TM, Ward NJ, et al. A structural model for binding of the serine-rich repeat adhesin GspB to host carbohydrate receptors. PLoS Pathog. 2011;7(7):e1002112.

16. Lizcano A, Sanchez CJ, Orihuela CJ. A role for glycosylated serine-rich repeat proteins in gram-positive bacterial pathogenesis. Mol Oral Microbiol. 2012;27(4):257–69.

17. Zhou M, Wu H. Glycosylation and biogenesis of a family of serine-rich bacterial adhesins. Microbiology. 2009;155(Pt 2):317–27.

18. Ramboarina S, Garnett JA, Zhou M, Li Y, Peng Z, Taylor JD, et al. Structural insights into serine-rich fimbriae from Gram-positive bacteria. J Biol Chem. 2010;285(42):32446–57.

19. Deng L, Bensing BA, Thamadilok S, Yu H, Lau K, Chen X, et al. Oral streptococci utilize a Siglec-like domain of serine-rich repeat adhesins to preferentially target platelet sialoglycans in human blood. PLoS Pathog. 2014;10(12):e1004540.

20. Bensing BA, Loukachevitch LV, Agarwal R, Yamakawa I, Luong K, Hadadianpour A, et al. Selectivity and engineering of the sialoglycan-binding spectrum in Siglec-like adhesins. bioRxiv. 2019:796912.

21. Bensing BA, Khedri Z, Deng L, Yu H, Prakobphol A, Fisher SJ, et al. Novel aspects of sialoglycan recognition by the Siglec-like domains of streptococcal SRR glycoproteins. Glycobiology. 2016;26(11):1222–34.

22. Urano-Tashiro Y, Takahashi Y, Oguchi R, Konishi K. Two Arginine Residues of Streptococcus gordonii Sialic Acid-Binding Adhesin Hsa Are Essential for Interaction to Host Cell Receptors. PLoS One. 2016;11(4):e0154098.

23. Xiong YQ, Bensing BA, Bayer AS, Chambers HF, Sullam PM. Role of the serine-rich surface glycoprotein GspB of Streptococcus gordonii in the pathogenesis of infective endocarditis. Microb Pathog. 2008;45(4):297–301.

24. Bensing BA, Li L, Yakovenko O, Wong M, Barnard KN, Iverson TM, et al. Recognition of specific sialoglycan structures by oral streptococci impacts the severity of endocardial infection. PLoS Pathog. 2019;15(6):e1007896.

25. Takahashi Y, Takashima E, Shimazu K, Yagishita H, Aoba T, Konishi K. Contribution of sialic acid-binding adhesin to pathogenesis of experimental endocarditis caused by Streptococcus gordonii DL1. Infect Immun. 2006;74(1):740–3.

26. Wong A, Grau MA, Singh AK, Woodiga SA, King SJ. Role of Neuraminidase-Producing Bacteria in Exposing Cryptic Carbohydrate Receptors for Streptococcus gordonii Adherence. Infect Immun. 2018;86(7).

27. Yang L, Connaris H, Potter JA, Taylor GL. Structural characterization of the carbohydrate-binding module of NanA sialidase, a pneumococcal virulence factor. BMC Struct Biol. 2015;15:15.

28. Bensing BA, Seepersaud R, Yen YT, Sullam PM. Selective transport by SecA2: an expanding family of customized motor proteins. Biochim Biophys Acta. 2014;1843(8):1674–86.

29. Singh AK, Pluvinage B, Higgins MA, Dalia AB, Woodiga SA, Flynn M, et al. Unravelling the multiple functions of the architecturally intricate Streptococcus pneumoniae beta-galactosidase, BgaA. PLoS Pathog. 2014;10(9):e1004364.

30. Marraffini LA, Dedent AC, Schneewind O. Sortases and the art of anchoring proteins to the envelopes of gram-positive bacteria. Microbiol Mol Biol Rev. 2006;70(1):192–221.

31. Schneewind O, Missiakas D. Sec-secretion and sortase-mediated anchoring of proteins in Gram-positive bacteria. Biochim Biophys Acta. 2014;1843(8):1687–97.

32. Navarre WW, Schneewind O. Proteolytic cleavage and cell wall anchoring at the LPXTG motif of surface proteins in gram-positive bacteria. Mol Microbiol. 1994;14(1):115–21.

33. McNab R, Holmes AR, Clarke JM, Tannock GW, Jenkinson HF. Cell surface polypeptide CshA mediates binding of Streptococcus gordonii to other oral bacteria and to immobilized fibronectin. Infect Immun. 1996;64(10):4204–10.

34. Holmes AR, McNab R, Jenkinson HF. Candida albicans binding to the oral bacterium Streptococcus gordonii involves multiple adhesin-receptor interactions. Infect Immun. 1996;64(11):4680–5.

35. McNab R, Jenkinson HF, Loach DM, Tannock GW. Cell-surface-associated polypeptides CshA and CshB of high molecular mass are colonization determinants in the oral bacterium Streptococcus gordonii. Mol Microbiol. 1994;14(4):743–54.

36. Back CR, Sztukowska MN, Till M, Lamont RJ, Jenkinson HF, Nobbs AH, et al. The Streptococcus gordonii Adhesin CshA Protein Binds Host Fibronectin via a Catch-Clamp Mechanism. J Biol Chem. 2017;292(5):1538–49.

37. Soding J, Biegert A, Lupas AN. The HHpred interactive server for protein homology detection and structure prediction. Nucleic Acids Res. 2005;33(Web Server issue):W244–8.

38. Christner M, Franke GC, Schommer NN, Wendt U, Wegert K, Pehle P, et al. The giant extracellular matrix-binding protein of Staphylococcus epidermidis mediates biofilm accumulation and attachment to fibronectin. Mol Microbiol. 2010;75(1):187–207.

39. Ronis A, Brockman K, Singh AK, Gaytan M, Wong A, McGrath S, et al. Streptococcus oralis subsp. dentisani produces mono-lateral serine rich repeat protein fibrils one of which contributes to saliva binding via sialic acid. Infect Immun. 2019.

40. Linke C, Siemens N, Oehmcke S, Radjainia M, Law RH, Whisstock JC, et al. The extracellular protein factor Epf from Streptococcus pyogenes is a cell surface adhesin that binds to cells through an N-terminal domain containing a carbohydrate-binding module. J Biol Chem. 2012;287(45):38178–89.

41. Crump KE, Bainbridge B, Brusko S, Turner LS, Ge X, Stone V, et al. The relationship of the lipoprotein SsaB, manganese and superoxide dismutase in Streptococcus sanguinis virulence for endocarditis. Mol Microbiol. 2014;92(6):1243–59.

42. Lowrance JH, Baddour LM, Simpson WA. The role of fibronectin binding in the rat model of experimental endocarditis caused by Streptococcus sanguis. J Clin Invest. 1990;86(1):7–13.

43. Que YA, Haefliger JA, Piroth L, Francois P, Widmer E, Entenza JM, et al. Fibrinogen and fibronectin binding cooperate for valve infection and invasion in Staphylococcus aureus experimental endocarditis. J Exp Med. 2005;201(10):1627–35.

44. Fitzgerald JR, Foster TJ, Cox D. The interaction of bacterial pathogens with platelets. Nat Rev Microbiol. 2006;4(6):445–57.

45. Khan R, Urban C, Rubin D, Segal-Maurer S. Subacute endocarditis caused by Gemella haemolysans and a review of the literature. Scand J Infect Dis. 2004;36(11-12):885–8.

46. Sadaune L, Roca F, Bordage M, Le Guillou V, Lesourd A, Michel A. Benefits of a Pre-Treatment Comprehensive Geriatric Assessment in a Rare Case of Gemella Haemolysans Endocarditis in an 86-Year-Old Patient and a Review of the Literature. Medicina (Kaunas). 2019;55(6).

47. Youssef D, Youssef I, Marroush TS, Sharma M. Gemella endocarditis: A case report and a review of the literature. Avicenna J Med. 2019;9(4):164–8.

48. Al-Tawfiq JA, Kiwan G, Murrar H. Granulicatella elegans native valve infective endocarditis: case report and review. Diagn Microbiol Infect Dis. 2007;57(4):439–41.

49. Ohara-Nemoto Y, Kishi K, Satho M, Tajika S, Sasaki M, Namioka A, et al. Infective endocarditis caused by Granulicatella elegans originating in the oral cavity. J Clin Microbiol. 2005;43(3):1405–7.

50. Padmaja K, Lakshmi V, Subramanian S, Neeraja M, Krishna SR, Satish OS. Infective endocarditis due to Granulicatella adiacens: a case report and review. J Infect Dev Ctries. 2014;8(4):548–50.

51. Patri S, Agrawal Y. Granulicatella elegans endocarditis: a diagnostic and therapeutic challenge. BMJ Case Rep. 2016;2016.

52. Ramnarain J, Yoon J, Runnegar N. Staphylococcus pasteuri infective endocarditis: A case report. IDCases. 2019;18:e00656.

53. Mitchell J. Streptococcus mitis: walking the line between commensalism and pathogenesis. Mol Oral Microbiol. 2011;26(2):89–98.

54. Kim SL, Gordon SM, Shrestha NK. Distribution of streptococcal groups causing infective endocarditis: a descriptive study. Diagn Microbiol Infect Dis. 2018;91(3):269–72.

55. Boratyn GM, Schaffer AA, Agarwala R, Altschul SF, Lipman DJ, Madden TL. Domain enhanced lookup time accelerated BLAST. Biol Direct. 2012;7:12.

56. Zheng W, Tan MF, Old LA, Paterson IC, Jakubovics NS, Choo SW. Distinct Biological Potential of Streptococcus gordonii and Streptococcus sanguinis Revealed by Comparative Genome Analysis. Sci Rep. 2017;7(1):2949.

57. Moschioni M, Pansegrau W, Barocchi MA. Adhesion determinants of the Streptococcus species. Microb Biotechnol. 2010;3(4):370–88.

58. Elliott D, Harrison E, Handley PS, Ford SK, Jaffray E, Mordan N, et al. Prevalence of Csh-like fibrillar surface proteins among mitis group oral streptococci. Oral Microbiol Immunol. 2003;18(2):114–20.

59. McNab R, Forbes H, Handley PS, Loach DM, Tannock GW, Jenkinson HF. Cell wall-anchored CshA polypeptide (259 kilodaltons) in Streptococcus gordonii forms surface fibrils that confer hydrophobic and adhesive properties. J Bacteriol. 1999;181(10):3087–95.

60. Connaris H, Crocker PR, Taylor GL. Enhancing the receptor affinity of the sialic acid-binding domain of Vibrio cholerae sialidase through multivalency. J Biol Chem. 2009;284(11):7339–51.

61. Goulas TK, Goulas AK, Tzortzis G, Gibson GR. Molecular cloning and comparative analysis of four beta-galactosidase genes from Bifidobacterium bifidum NCIMB41171. Appl Microbiol Biotechnol. 2007;76(6):1365–72.

62. Chitayat S, Adams JJ, Smith SP. NMR assignment of backbone and side chain resonances for a dockerin-containing C-terminal fragment of the putative mu-toxin from Clostridium perfringens. Biomol NMR Assign. 2007;1(1):13–5.

63. Chitayat S, Adams JJ, Furness HS, Bayer EA, Smith SP. The solution structure of the C-terminal modular pair from Clostridium perfringens mu-toxin reveals a noncellulosomal dockerin module. J Mol Biol. 2008;381(5):1202–12.

64. Fujita K, Oura F, Nagamine N, Katayama T, Hiratake J, Sakata K, et al. Identification and molecular cloning of a novel glycoside hydrolase family of core 1 type O-glycan-specific endo-alpha-N-acetylgalactosaminidase from Bifidobacterium longum. J Biol Chem. 2005;280(45):37415–22.

65. Ashida H, Miyake A, Kiyohara M, Wada J, Yoshida E, Kumagai H, et al. Two distinct alpha-L-fucosidases from Bifidobacterium bifidum are essential for the utilization of fucosylated milk oligosaccharides and glycoconjugates. Glycobiology. 2009;19(9):1010–7.

66. Trastoy B, Lomino JV, Pierce BG, Carter LG, Gunther S, Giddens JP, et al. Crystal structure of Streptococcus pyogenes EndoS, an immunomodulatory endoglycosidase specific for human IgG antibodies. Proc Natl Acad Sci U S A. 2014;111(18):6714–9.

67. Dixon EV, Claridge JK, Harvey DJ, Baruah K, Yu X, Vesiljevic S, et al. Fragments of bacterial endoglycosidase s and immunoglobulin g reveal subdomains of each that contribute to deglycosylation. J Biol Chem. 2014;289(20):13876–89.

68. Cheng AG, Missiakas D, Schneewind O. The giant protein Ebh is a determinant of Staphylococcus aureus cell size and complement resistance. J Bacteriol. 2014;196(5):971–81.

69. Clarke SR, Harris LG, Richards RG, Foster SJ. Analysis of Ebh, a 1.1-megadalton cell wall-associated fibronectin-binding protein of Staphylococcus aureus. Infect Immun. 2002;70(12):6680–7.

70. Tanaka Y, Sakamoto S, Kuroda M, Goda S, Gao YG, Tsumoto K, et al. A helical string of alternately connected three-helix bundles for the cell wall-associated adhesion protein Ebh from Staphylococcus aureus. Structure. 2008;16(3):488–96.

71. Lin IH, Hsu MT, Chang CH. Protein domain repetition is enriched in Streptococcal cell-surface proteins. Genomics. 2012;100(6):370–9.

72. Schroeder K, Jularic M, Horsburgh SM, Hirschhausen N, Neumann C, Bertling A, et al. Molecular characterization of a novel Staphylococcus aureus surface protein (SasC) involved in cell aggregation and biofilm accumulation. PLoS One. 2009;4(10):e7567.

73. Velez MP, Petrova MI, Lebeer S, Verhoeven TL, Claes I, Lambrichts I, et al. Characterization of MabA, a modulator of Lactobacillus rhamnosus GG adhesion and biofilm formation. FEMS Immunol Med Microbiol. 2010;59(3):386–98.

74. Shivshankar P, Sanchez C, Rose LF, Orihuela CJ. The Streptococcus pneumoniae adhesin PsrP binds to Keratin 10 on lung cells. Mol Microbiol. 2009;73(4):663–79.

75. Takahashi Y, Yajima A, Cisar JO, Konishi K. Functional analysis of the Streptococcus gordonii DL1 sialic acid-binding adhesin and its essential role in bacterial binding to platelets. Infect Immun. 2004;72(7):3876–82.

76. Wu H, Zeng M, Fives-Taylor P. The glycan moieties and the N-terminal polypeptide backbone of a fimbria-associated adhesin, Fap1, play distinct roles in the biofilm development of Streptococcus parasanguinis. Infect Immun. 2007;75(5):2181–8.

77. Handley PS, Correia FF, Russell K, Rosan B, DiRienzo JM. Association of a novel high molecular weight, serine-rich protein (SrpA) with fibril-mediated adhesion of the oral biofilm bacterium Streptococcus cristatus. Oral Microbiol Immunol. 2005;20(3):131–40.

78. Seo HS, Xiong YQ, Sullam PM. Role of the serine-rich surface glycoprotein Srr1 of Streptococcus agalactiae in the pathogenesis of infective endocarditis. PLoS One. 2013;8(5):e64204.

79. Kilian M, Tettelin H. Identification of Virulence-Associated Properties by Comparative Genome Analysis of Streptococcus pneumoniae, S. pseudopneumoniae, S. mitis, Three S. oralis Subspecies, and S. infantis. mBio. 2019;10(5).

80. Reichmann P, Konig A, Linares J, Alcaide F, Tenover FC, McDougal L, et al. A global gene pool for high-level cephalosporin resistance in commensal Streptococcus species and Streptococcus pneumoniae. J Infect Dis. 1997;176(4):1001–12.

81. Granok AB, Parsonage D, Ross RP, Caparon MG. The RofA binding site in Streptococcus pyogenes is utilized in multiple transcriptional pathways. J Bacteriol. 2000;182(6):1529–40.

82. Bishop CJ, Aanensen DM, Jordan GE, Kilian M, Hanage WP, Spratt BG. Assigning strains to bacterial species via the internet. BMC Biol. 2009;7:3.

83. Jensen A, Scholz CFP, Kilian M. Re-evaluation of the taxonomy of the Mitis group of the genus Streptococcus based on whole genome phylogenetic analyses, and proposed reclassification of Streptococcus dentisani as Streptococcus oralis subsp. dentisani comb. nov., Streptococcus tigurinus as Streptococcus oralis subsp. tigurinus comb. nov., and Streptococcus oligofermentans as a later synonym of Streptococcus cristatus. Int J Syst Evol Microbiol. 2016;66(11):4803–20.

84. Tamura K, Stecher G, Peterson D, Filipski A, Kumar S. MEGA6: Molecular Evolutionary Genetics Analysis version 6.0. Mol Biol Evol. 2013;30(12):2725–9.

85. Fujiwara T, Hoshino T, Ooshima T, Sobue S, Hamada S. Purification, characterization, and molecular analysis of the gene encoding glucosyltransferase from Streptococcus oralis. Infect Immun. 2000;68(5):2475–83.

86. Koren S, Walenz BP, Berlin K, Miller JR, Bergman NH, Phillippy AM. Canu: scalable and accurate long-read assembly via adaptive k-mer weighting and repeat separation. Genome Res. 2017;27(5):722–36.

87. Simao FA, Waterhouse RM, Ioannidis P, Kriventseva EV, Zdobnov EM. BUSCO: assessing genome assembly and annotation completeness with single-copy orthologs. Bioinformatics. 2015;31(19):3210–2.

88. Seemann T. Prokka: rapid prokaryotic genome annotation. Bioinformatics. 2014;30(14):2068–9.

89. Almagro Armenteros JJ, Tsirigos KD, Sonderby CK, Petersen TN, Winther O, Brunak S, et al. SignalP 5.0 improves signal peptide predictions using deep neural networks. Nat Biotechnol. 2019;37(4):420–3.

90. Nawrocki EP, Eddy SR. Infernal 1.1: 100-fold faster RNA homology searches. Bioinformatics. 2013;29(22):2933–5.

91. McLaughlin RE, Ferretti JJ. Electrotransformation of Streptococci. Methods Mol Biol. 1995;47:185–93.

92. Alam S, Brailsford SR, Whiley RA, Beighton D. PCR-Based methods for genotyping viridans group streptococci. J Clin Microbiol. 1999;37(9):2772–6.

93. Wiedemann C, Bellstedt P, Gorlach M. CAPITO--a web server-based analysis and plotting tool for circular dichroism data. Bioinformatics. 2013;29(14):1750–7.

94. Rueden CT, Schindelin J, Hiner MC, DeZonia BE, Walter AE, Arena ET, et al. ImageJ2: ImageJ for the next generation of scientific image data. BMC Bioinformatics. 2017;18(1):529.

95. Waterhouse A, Bertoni M, Bienert S, Studer G, Tauriello G, Gumienny R, et al. SWISS-MODEL: homology modelling of protein structures and complexes. Nucleic Acids Res. 2018;46(W1):W296–W303.

96. Madeira F, Park YM, Lee J, Buso N, Gur T, Madhusoodanan N, et al. The EMBL-EBI search and sequence analysis tools APIs in 2019. Nucleic Acids Res. 2019;47(W1):W636–W41.

