## Supplementary material for "A novel sialic acid-binding adhesin present in multiple species contributes to the pathogenesis of Infective endocarditis": Table S1

| **Gene ID** | **GenBank ID** | **Length (aa)** | **BLAST hits** | **Pfam domains** |
| --- | --- | --- | --- | --- |
| **Present in IE12 and IE18** |  |  |  |  |
| IE12_1004 (IE18_1293) | HRE59_04885  (HRJ32_06265) | 2143 | S8 family serine peptidase | Inhibitor_I9 (PF05922)  Peptidase_S8 (PF00082)  fn3_5 (PF06280) |
| IE12_1078 (IE18_1187) | HRE59_05235  (HRJ32_05765) | 459 | VWA domain-containing protein | VWA (PF00092) |
| IE12_1764 (IE18_0535) | HRE59_08575  (HRJ32_02590) | 3146 | DUF1542 domain-containing protein | DUF1542 (PF07564) x 31 |
| IE12_1765 (IE18_0534) | HRE59_08580 (HRJ32_02585) | 1929 | Immunoglobulin A1 protease precursor | G5 (PF07501) x 2  Peptidase_M26_N (PF05342)  Peptidase_M26_C (PF07580) |
| **Present only in IE18** |  |  |  |  |
| IE18_0557 | HRJ32_02705 | 3097 | YSIRK signal domain/LPXTG anchor domain surface protein | CshA_NR2 (PF18651) |
| IE18_1262 | HRJ32_06115 | 274 | LPXTG cell wall anchor domain-containing protein | - |
| IE18_1284 | HRJ32_06220 | 976 | Major cell-surface adhesin PAc precursor | Adhesin_P1_N (PF18652)  AgI_II_C2 (PF17998)  Antigen_C (PF16364) |
| IE18_2031 | HRJ32_09925 | 836 | SEC10/PgrA surface exclusion domain-containing protein | - |

fn3_5: Fibronectin type III domain

VWA: Von Willebrand factor type A

DUF: Domain of Unknown Function

Peptidase_M26_N: M26 IgA1-specific Metallo-endopeptidase N-terminal region

Peptidase_M26_C: M26 IgA1-specific Metallo-endopeptidase C-terminal region

Adhesin_P1_N: Adhesin P1 N-terminal domain

AgI_II_C2: Cell surface antigen I/II C2 terminal domain

Antigen_C: Cell surface antigen C-terminus

CshA_NR2: Surface adhesin CshA non-repetitive domain 2
