## Supplementary figures and images for "A novel sialic acid-binding adhesin present in multiple species contributes to the pathogenesis of Infective endocarditis"

### Figure S1

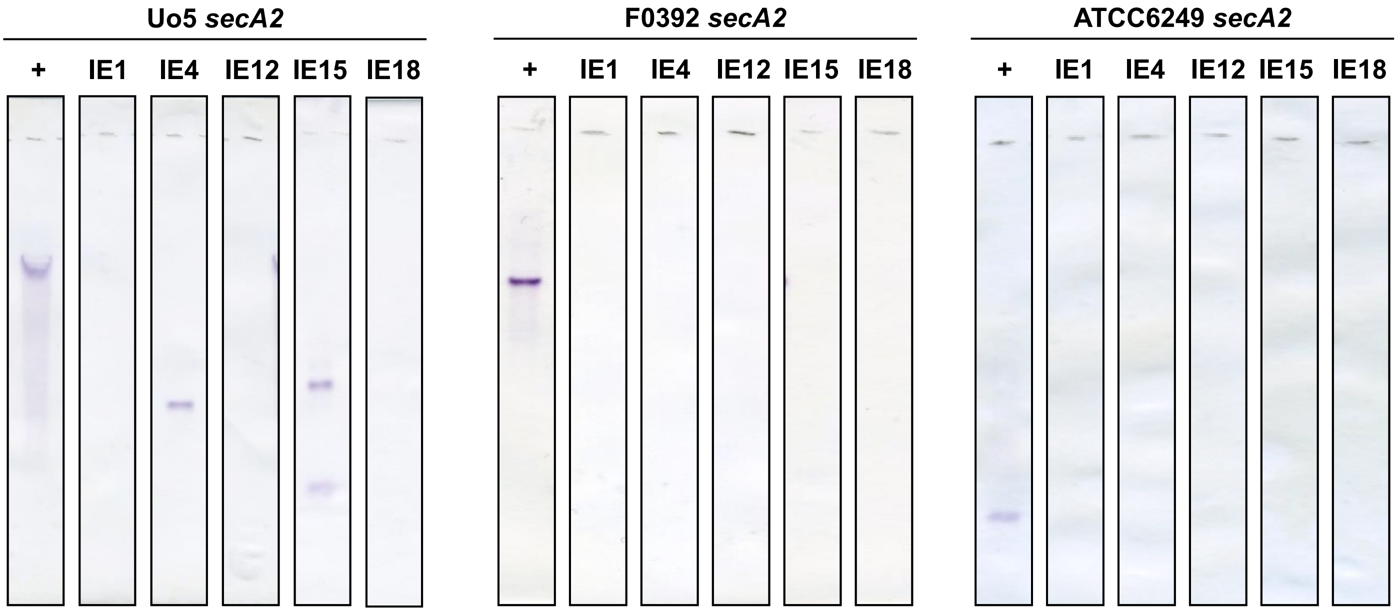

### Figure S2

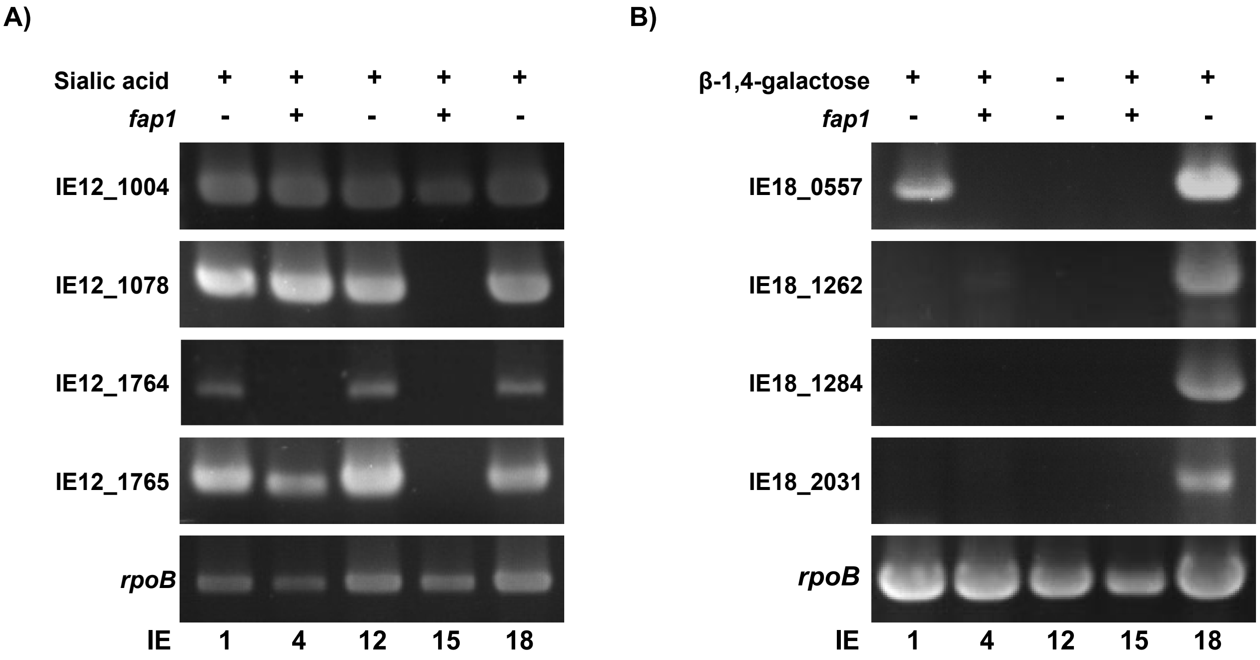

### Figure S3

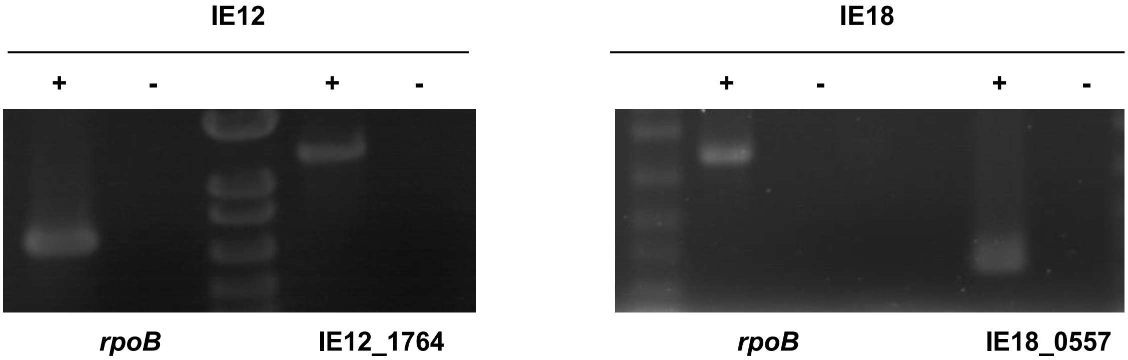

### Figure S4

**IE18**

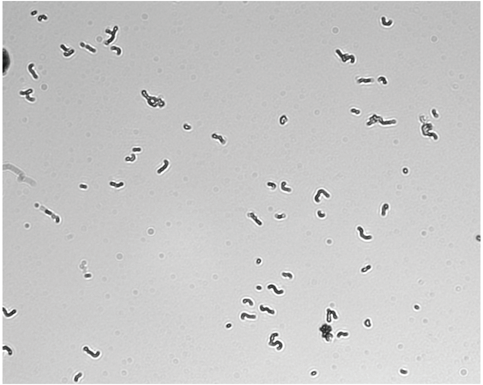

**IE18 Δ*csh-like***

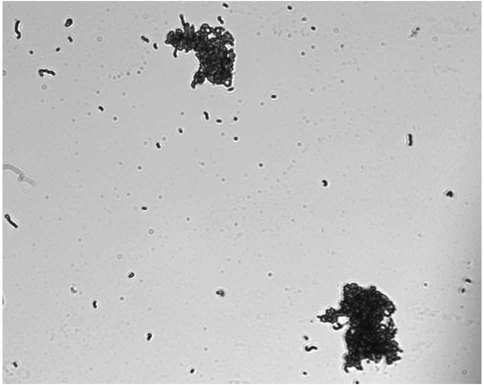

### Figure S5

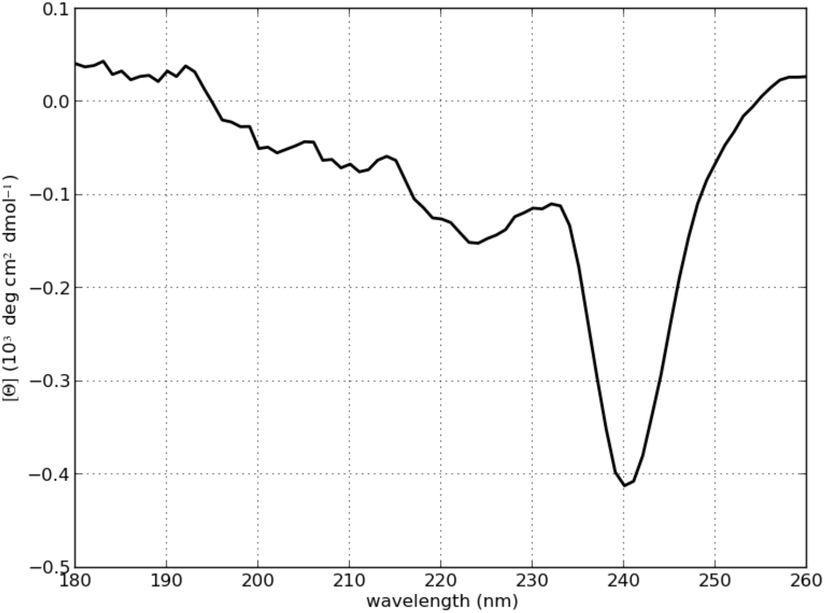

### Figure S6

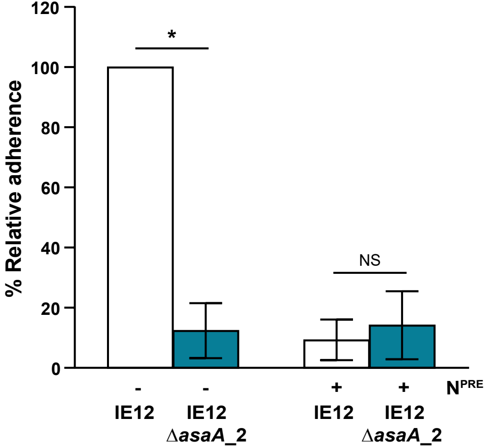

### Figure S7

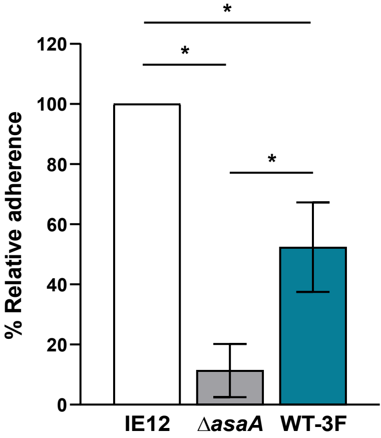

### Figure S8

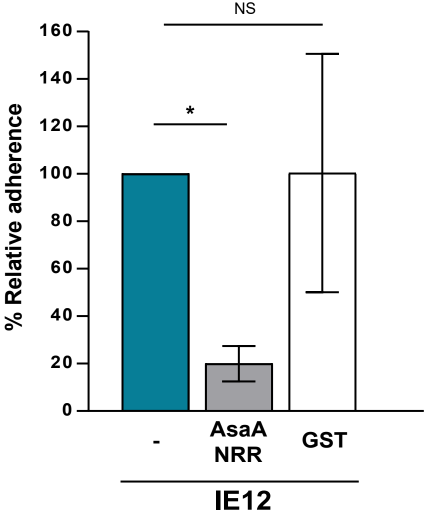

### Figure S9

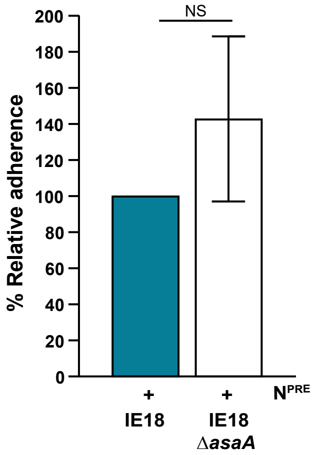
